# *Bacillus subtilis* derived lipopeptides disrupt quorum sensing and biofilm assembly in *Staphylococcus aureus*

**DOI:** 10.1101/2023.08.24.554662

**Authors:** Kyle R. Leistikow, Daniel S. May, Won Se Suh, Gabriel Vargas Asensio, Cameron R. Currie, Krassimira R. Hristova

## Abstract

Multidrug-resistant *Staphylococcus aureus* is one of the most clinically important pathogens in the world with infections leading to high rates of morbidity and mortality in both humans and animals. *S. aureus’* ability to form biofilm protects individual cells from antibiotics and promotes the transfer of antibiotic resistance genes. Therefore, new strategies aimed to inhibit biofilm growth and disassemble mature biofilms are urgently needed. Probiotic species, namely *Bacillus subtilis,* are gaining interest as a potential therapeutic against *S. aureus* for their ability to reduce *S. aureus* colonization and virulence. Here, we collected and screened 1123 *Bacillus* strains obtained from a variety of agricultural environments in search of isolates with strong antibiofilm activity against clinical multi-drug resistant *S. aureus.* We selected a single strain, *B. subtilis* 6D1, based on its ability to inhibit biofilm growth, disassemble mature biofilm, and improve antibiotic sensitivity of *S. aureus* biofilms through an Agr quorum sensing interference mechanism. Biochemical and molecular networking analysis of an active organic fraction revealed multiple surfactin isoforms and an uncharacterized compound were both driving this antibiofilm activity. Furthermore, when compared against commercial HPLC grade surfactin obtained from *B. subtilis,* this active fraction inhibited biofilm formation against all four *S. aureus* Agr backgrounds and prevented *S. aureus*-induced cytotoxicity when applied to HT29 human intestinal cell lines better than the commercial standard. Our results demonstrate the mixture of compounds produced by *B. subtilis* 6D1 can mitigate *S. aureus* virulence through multiple mechanisms.

**Contribution to the Field:** The biofilm formation capability of bacterial pathogens, such as *Staphylococcus aureus*, increases these microorganisms’ virulence potential and decreases the efficacy of common antibiotic regiments. Probiotics possess a variety of strain-specific strategies to reduce biofilm formation in competing organisms, however, the mechanisms and compounds responsible for these phenomena often go uncharacterized. In this study, we identified a mixture of small probiotic-derived peptides capable of Agr quorum sensing interference as one of the mechanisms driving antibiofilm activity against *S. aureus.* This collection of peptides also improved antibiotic killing and protected human gut epithelial cells from *S. aureus*-induced toxicity by stimulating an adaptive immune response. We conclude that purposeful strain screening and selection efforts can be used to identify unique probiotic strains that possess specially desired mechanisms of action. This information can be used to further improve our understanding of the ways in which probiotic and probiotic-derived compounds can be applied to prevent bacterial infections in clinical and agricultural settings.

## Introduction

Antibiotic resistant *Staphylococcus aureus* is a leading cause of pneumonia, sepsis, endocarditis, and soft tissue infections. In addition to its constantly evolving resistance to front-line antibiotics^1^, *S. aureus* infections are made more difficult to treat due to its biofilm formation abilities. A variety of synthetic compounds have been investigated for their ability to reduce *S. aureus* biofilm growth^2–4^, however, very little research has investigated the effects of probiotics and probiotic derived small molecules on pathogenic biofilms. Probiotics possess a variety of strain specific strategies to reduce pathogen growth, virulence, and/or biofilm formation and have shown extraordinary potential to prevent and treat a variety of diseases^5,6^, including *S. aureus* infections^7^. The majority of probiotic research has sought to identify strains with broad bactericidal activity^8^, however, recent evidence suggests some strains can reduce disease outcomes by inhibiting pathogen quorum sensing, a population-dependent bacterial communication system that modulates gene expression and lifestyle selection. Quorum sensing inhibitors do not kill bacteria; instead, they interfere with communication systems needed to form biofilms and produce virulence factors. Quorum sensing interference (QSI) has also been proposed as a method to improve antibiotic killing effects^9,10^, making it an intriguing mechanism with immediate therapeutic potential.

In *S. aureus*, quorum sensing (QS) is controlled through the secretion and self-recognition of post-translationally modified autoinducing peptides. Specifically, the agrBDCA operon, integral to a functioning Agr QS system, allows *S. aureus* to coordinate the formation and disassembly of biofilm and is required for approximately 90% of *S. aureus* infections^11^. The Agr QS system is conserved among *S. aureus* strains, but variations within AgrD and the C-terminus of AgrB result in four different mature autoinducing peptides (AIPs) that are uniquely recognized by four differently configured AgrC receptors^12^. Each of the four *S. aureus* Agr groups has a different biological consequence: Agr-I is linked to enterotoxin disease; Agr-II is linked to early vancomycin-resistance and endocarditis; Agr-III is linked to endocarditis and menstrual toxic shock syndrome, and Agr-IV is linked to exfoliative disease^13,14^. Despite these differences, the Agr QS system in each group is responsible for regulating virulence factor gene expression^15^ and facilitating biofilm formation and dispersal^16,17^. When the Agr QS system is turned on, *S. aureus* maintains a planktonic lifestyle. Conversely, when the Agr QS system is turned off, *S. aureus* forms biofilm. Importantly, this phenomena can be impeded by competitive Agr signaling interference^18^. This occurs when peptides produced by different species bind to AgrC and successfully initiate the quorum sensing pathways in neighboring cells^19,20^. Therefore, it is possible that certain probiotic strains produce peptides that reduce *S. aureus* biofilm formation through an activation of the Agr QS system^21^.

One such probiotic species, *Bacillus subtilis*, is gaining interest as a potential therapeutic against *S. aureus*. Though the potential health benefits associated with *B. subtilis* are well documented, the reproducibility of these interventions largely depends on the *B. subtilis* strain used^22,23^. *B. subtilis* strains are remarkably diverse and shaped both by their environment and their ability to acquire genes from closely related species^24,25^; therefore, due to *B. subtilis’* diverse ecological range, it is perhaps not surprising that strain specific genetic elements have evolved to help this species compete in a variety of environments^26,27^*. B. subtilis* excretes a variety of compounds^28^ that inhibit *S. aureus* biofilm growth^29^, however since the majority of these experiments use crude *Bacillus* cell-free extracts, the precise compounds driving these phenomenon often go unidentified^21^.

Here, we screened a library of *Bacillus* strains sourced from a variety of contaminated dairy environments for antibiofilm activity against *S. aureus*. We hypothesized that potential probiotic strains obtained from these environments had likely co-evolved to compete with a variety of dairy pathogens, namely *S. aureus*, the primary etiological agent responsible for bovine mastitis^30,31^. Therefore, we investigated how *Bacillus subtilis* inhibits *S. aureus* biofilm growth, identified the compound(s) driving this activity, and then assessed these compounds’ protective effects against *S. aureus* infection in eukaryotic cell culture experiments. Investigating probiotic-mediated mechanisms that reduce virulence and improve antibiotic efficacy, and identifying the *B. subtilis*-derived products responsible, is essential to understand how probiotic microorganisms can be deployed to combat *S. aureus* infections and help to reduce the use of antibiotics intended to treat these multi-drug resistant pathogens.

## Methods

### Strains and Chemicals Used in this Study

Please see supplementary Table S1 for a list of strains used in this study Surfactin from *Bacillus subtilis* >98% (CAS: 24730-31-2) (Lot:0000134293) (Source:0000128758) (Sigma-Aldrich, USA) Fengycin from *Bacillus subtilis* ≥ 90% (CAS: 102577-03-7) (Source: SLCK5954) (Sigma-Aldrich, USA) Lysostaphin (CAS:9011-93-2) (Lot:109M4010V) (Sigma-Aldrich, USA) Dimethyl Sulfoxide (CAS:67-68-5) (Lot: 184176) (Fisher Chemical, USA)

### *Bacillus* Candidate Screening

To attain presumptive *Bacillus* isolates, 50 environmental samples comprised of bovine feces and milk, and manure contaminated river sediment and soil were collected across Kewaunee County, Wisconsin – home to 16 concentrated animal feeding operations and more than 200 smaller dairy farms. One gram per sample was applied to 9mL 0.1% peptone, heated at 72°C for 20 minutes to induce *Bacillus* sporulation, and plated aerobically on trypticase soy agar (TSA) plates. To ensure only a single strain was obtained, a total of 1123 presumptive *Bacillus* colonies were individually picked with sterile toothpicks and inoculated into fresh square TSA plates in a 6 x 6 grid. To reduce overgrowth from any fastidious strains, plates were grown at room temperature for 48 hours. After incubation, each strain was sub-cultured on agar plates to ensure pure culture isolation and subsequently in 500μL trypticase soy broth (TSB) in 96-well deep v-well reservoirs (Thermo Scientific, USA). All cultures were grown for 24 hours at 37°C and used immediately for antimicrobial screening assays.

All *Bacillus* strains were assessed for antagonistic activity against methicillin resistant and methicillin susceptible clinical and type strains of *S. aureus* using drop diffusion overlay and cross streak assays^32^. For drop diffusion assays, all 1123 presumptive *Bacillus* isolates were grown for 24 hours in TSB at 37°C and 5μl were spotted onto square petri plates (VWR International, USA) with TSA media in a 6 x 6 grid fashion and allowed to dry. Seven mL of soft TSA (0.75% agar) harboring 150μL of exponential phase *S. aureus* cultures was overlaid onto the TSA plate and allowed to dry. Plates were then inverted and incubated at 37°C for 18 hours before measuring clearing zones indicative of antagonistic activity. A total of 75 strains exhibiting inhibition against both *S. aureus* strains were subsequently assayed using cross streak methods. Briefly, 100μL of each presumptive *Bacillus* culture was inoculated into 10mL TSB and incubated overnight at 37°C. A loop full of each culture was struck vertically down the center of a square TSA plate and incubated for an additional 16 hours at 37°C. The next day, nine different methicillin susceptible and methicillin resistant clinical and type strains including the two used previously for drop diffusion assays, were struck perpendicularly towards the *Bacillus* streak, allowed to dry, and incubated at 37°C overnight prior to measuring zones of inhibition.

*Bacillus* strains (n=46) exhibiting broad antagonistic activity against *S. aureus* in both assays were individually cultured in 125mL baffled shake flasks and incubated at 37°C shaking at 230 RPM for 48 hours prior to harvesting cell-free extracts for use in subsequent biofilm and planktonic inhibition assays. After incubation, cultures were centrifuged at 4°C at 10,500 x g for 20 minutes. Cell-free extract (CFE) was harvested and filter-sterilized using a 0.22μm filter (Corning, USA). CFEs were flash frozen in liquid nitrogen and stored at −80°C for future use. A total of 39 clinical methicillin susceptible and methicillin resistant *S. aureus* strains were used to assess antibiofilm activity of *Bacillus* CFEs. A list of all Staphylococcal strains used throughout this work is outlined in Supplementary Table 1. *Bacillus* strains (n=16) exhibiting broad antimicrobial potential in a planktonic or biofilm environment were assayed for relevant pathogenic *Bacillus* toxin genes^33^ prior to sequencing the 16S rRNA gene region for taxonomic classification. Among these 16 candidates, a single strain, *B. subtilis* 6D1, exhibited stronger antibiofilm activity without affecting planktonic growth and thus was selected for more detailed investigations

### Whole Genome Sequencing & Bioinformatics Analysis

Whole genome sequencing and assembly was performed on *B. subtilis* 6D1 to conduct comparative genomic studies and to mine for potentially unique genetic traits. Briefly, *Bacillus* DNA was isolated using DNeasy Blood and Tissue Kit (Qiagen, Germany) according to the manufacturer’s instructions. To improve sequencing accuracy at the nucleotide level, both Illumina (400Mbs) and Nanopore (300Mbs) sequencing was performed at SeqCenter (Pittsburgh, USA). Quality control and adapter trimming was performed with bcl2fastq^34^ and porechop^35^ for Illumina and Nanopore sequencing, respectively. Hybrid read assembly was performed *de novo* with Unicycler^36^, and assembly statistics were recorded with QUAST^37^. The taxonomy classification of the *B. subtilis* 6D1 genome assembly was confirmed using GTDB-tk (Gene Bank Accession # CP129123-CP129124). Assemblies were annotated with Prokka^38^, CARD^39^, Phigaro^40^, and VirSorter2^41^. Genome maps were created using Proksee^38^. Assemblies were also analyzed in AntiSmash 6.0 to search for unique biosynthetic gene clusters (BGCs)^42^. Unique BGCs were further compared to reference BGCs using Clinker^43^. Phylogenetic analysis comparing *B. subtilis* 6D1 to 98 similarly identified Refseq genomes using 500 random single copy protein coding genes was performed using Minhash similar genome finder and codon tree builder pipelines found in the Bacterial and Viral Bioinformatics Resource Center (BV-BRC)^44^.

All available genomes of *Bacillus subtilis* on the IMG database were downloaded and phylogenomic analysis was performed using Anvi’o 6.2^45^. Specifically, we annotated the single copy core genes in all the genomes, concatenated a set of 71 core genes and aligned them in Anvi’o. The phylogenomic tree was constructed using FastTree 2.1.10^46^ and edited using iTOL^47^. Pangenome analyses utilized only genomes that were most closely related to *B. subtilis* 6D1 in the phylogenomic tree (purple clade in Figure 1C). Pangenome analysis was performed in Anvi’o as follows: a genome storage database was generated comprised of the *B. subtilis* 6D1 assembly and each of the closely related genomes; the pangenome analysis was run using the following parameters: minbit=05 and a mcl-inflation= 0.9 for the clustering. Diamond was used to compare the genes in sensitive mode^48^. The pangenome was visualized, inspected, and edited in Anvi’o. Annotation of unique *B. subtilis* 6D1 genes was performed in EggNOG-mapper v2.1.12.

**Figure 1.**
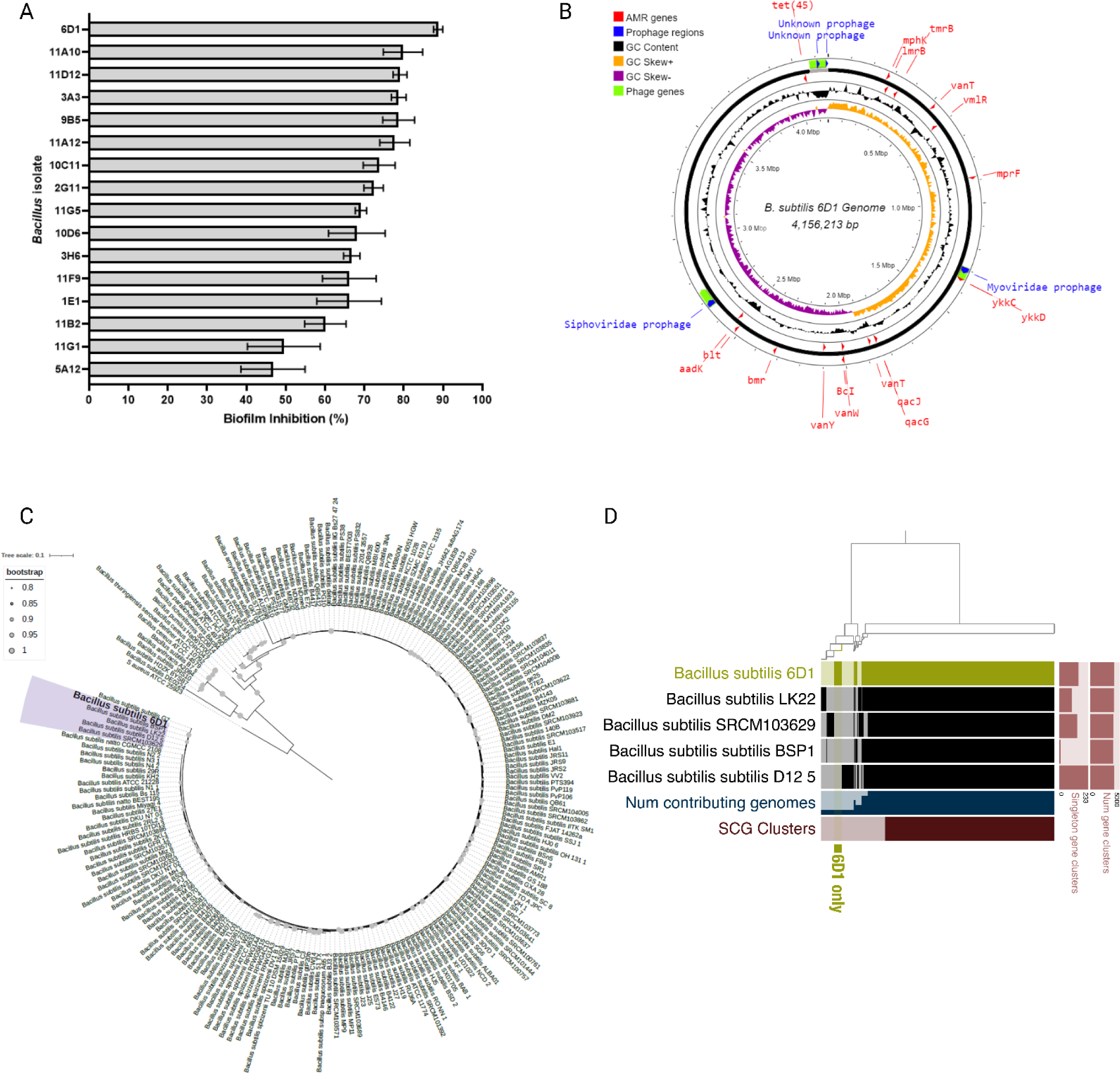
*B. subtilis* 6D1 exhibits antibiofilm activity against *S. aureus* and harbors unique genetic traits not observed in closely related *B. subtilis* strains. **(A)** Antibiofilm activity of 16 candidate *Bacillus* cell-free extracts against *S. aureus* 29213. **(B)** The assembled *B. subtilis* 6D1 genome **(C)** *B. subtilis* 6D1 shares >0.95% identity with a group of *B. subtilis* genomes (purple), however **(D)** pangenome analysis identified 156 unique genes in *B. subtilis* 6D1 that were not detected in any strain from this closely related group. Singleton gene clusters identified range from 0-233, number of gene clusters identified range from 0-5000.

### Competition Assays

For planktonic competitions, 250μL (5 x 10^7^ CFU) of exponential phase *B. subtilis* 6D1 and *S. aureus* ATCC 29213 were added to 25mL TSB in a baffled shake flask and incubated at 37°C shaking at 200 RPM. Biofilm competitions utilized 7 mm polystyrene beads (Polysciences, Inc, Warrington, PA) as a surface for cell attachment, biofilm growth, and biofilm dispersal^49^. For biofilm competitions, equal concentrations of *B. subtilis* 6D1 and *S. aureus* populations (∼1 x 10^5^ – 1 x 10^6^) were added to a glass test tube containing 5mL of 1.5% TSB +0.3% glucose media^50^ and a single polystyrene bead^51,52^. After 24 hours incubation at 37°C in a roller drum, the bead was placed in 1mL PBS and sonicated with a handheld Qsonica model CL-188 instrument (QSonica, USA) for 10 seconds at 60Hz to remove attached cells. Detached cells were serially diluted and plated on TSA for CFU enumeration. For all competitions, CFU counts were obtained for both bacterial species at time point 0 and after 24 hours incubation. Strain fitness and selection rate were calculated as previously described by Travisano and Lenski^53^.

### Biofilm Inhibition & Disruption Assays

*Bacillus* CFEs were applied at varying concentrations to Nunc™ treated 96 well microplates (Thermo Scientific, USA) containing 1.5% TSB + 0.3% glucose media^50^. Test wells were seeded with 5 x 10^5^ CFU/mL of *S. aureus* and incubated statically at 37°C. After 24 hours, media was removed, and biofilms were gently washed three times with PBS, fixed for 20 minutes at 55°C, and stained with 0.1% crystal violet (CV) for 15 minutes. After 15 minutes, CV was removed, and wells were washed twice with PBS and allowed to dry. CV was solubilized with 30% acetic acid. Biofilm inhibition was measured spectrophotometrically relative to untreated control wells at OD_595_. To assess biofilm disruption, *Bacillus* CFE (2.5 – 20% v/v) was mixed with PBS and applied to *S. aureus* 24-hour mature biofilms and incubated at 37°C shaking at 100 RPM for 2 hours. To account for any mechanical disruption not driven by CFE, biofilms treated only with PBS were used as a negative control. Residual biofilms were fixed for 20 minutes at 55°C and quantified by crystal violet staining as outlined above. Assays were performed in quadruplicate and averaged across three independent experiments. Biofilm inhibition and removal were confirmed via CFU plating after the initial PBS wash but prior to any CV staining.

### Antibiotic Synergy Biofilm Inhibition Assays

*Bacillus* CFEs exhibiting antibiofilm potential were applied in conjunction with ampicillin, gentamicin, and trimethoprim. Antibiotic concentrations were based on preliminary biofilm and planktonic minimum inhibitory concentrations and recommended CLSI breakpoints^54^. Checkerboard assays and fractional inhibitory concentration indices were used to assess whether coadministration of antibiotic and *Bacillus* CFE was synergistic(≤0.5), additive (0.5-1.0), or indifferent (1-4)^32^. Briefly, 5 x 10^5^ CFU/mL *S. aureus* ATCC 29213 was inoculated in polystyrene 96-well plates and treated with either 10% v/v *Bacillus* CFE, antibiotic (either gentamicin, ampicillin, or trimethoprim) or a combined antibiotic + *Bacillus* CFE treatment. Each antibiotic or antibiotic + *Bacillus* CFE combination was added at time point 0 and biofilm growth was measured via CV staining after 24h at 37°C incubation.

### Gene Expression

RNA was isolated from 24 hour broth cultures seeded with 5 x 10^5^ CFU/mL *S. aureus* ATCC 29213 co-inoculated with either *B. subtilis* 6D1 cell-free extract (10% v/v) or 5 x 10^5^ CFU/mL *B. subtilis* 6D1 cells (1:1) using a Qiagen RNeasy Mini kit adapted to include a modified lysis step with lysostaphin^55^. Prior to RNA extraction, each treated culture was mixed 1:2 with RNA Protect (Qiagen, Germany), vortexed for 10 seconds, and incubated at ambient temperature for 10 minutes. 200μL of lysis buffer (30mM Tris, 1mM EDTA, 20mg/mL lysozyme, 100μg/mL lysostaphin, pH= 8) and 20uL proteinase K were applied to each culture and incubated at ambient temperature on a rotary shaker (Fisher Scientific, USA) for 45 minutes. RNA quality and quantity were determined with a Qubit spectrophotometer (Thermo Scientific, USA). cDNA was obtained using the Qiagen Quantitect reverse transcriptase kit according to the manufacturer’s instructions (Qiagen, Germany). qPCR conditions were optimized and evaluated on a BioRad CFX96 real time instrument (Bio-Rad Laboratories, USA) based on previous work^3^. RT-qPCR reaction volumes (20μL) were comprised of 10ng cDNA, 300nM forward and reverse primers, and SYBR green master mix (Bio-Rad Laboratories, USA). The real-time cycling conditions were as follows: 95°C for 10 min, followed by 40 cycles of 95°C for 15 seconds and between 58.5-60°C (based on primer melting temperature) for 1 minute. Gene expression was quantified relative to the *rpoB* gene^3,56^ and calculated using the Livak method^57^. RNA expression was determined using three independent bacterial cultures tested in triplicate. Gene primers and their associated melting temperatures are listed in supplementary table 2.

### Confocal Laser Scanning Microscopy

To assess impact of *B. subtilis* 6D1 compounds on *S. aureus* biofilm formation, ibidi 8-μwell coated plates (ibidi, Germany) containing 400 μl 1.5% TSB + 0.3% glucose were used to grow and subsequently stain residual biofilms. Bacterial viability staining was performed using the Live/Dead BacLight Bacterial Viability Kit (Thermo Scientific, USA) according to manufacturer’s instructions. Briefly, both SYTO-9 and propidium iodide (PI) stains were thawed away from light and mixed with sterile distilled water at a final concentration of 0.1% for each dye. This mixture was equilibrated at ambient temperature for 10 minutes, and 200μL of the resulting mixture was applied to each well and incubated at ambient temperature for 15 minutes in the dark. Prior to imaging, stain solution was removed, and biofilms were washed twice with sterile PBS. Biofilms were observed using a 40X objective and measured via confocal laser scanning microscopy (CLSM) on a Nikon Eclipse Ti instrument. Five pre-determined fields of view were imaged per well on three independent wells per biofilm culture condition. Images and mean fluorescence intensity were analyzed in Imaris (version 9.9).

### *Bacillus* CFE Preparation for LC-MS/MS Analysis

To extract and separate lipopeptides from crude *Bacillus* CFE, liquid-liquid extractions using ethyl acetate were performed as previously described^58–60^. Briefly, *Bacillus* strains were cultured in 500mL TSB in 2L baffled shake flasks for 48 hours and centrifuged (4300 RPM, 30 min) at 4°C. Three liters of culture supernatant were mixed in a 1:1 volume ratio (v/v) with ethyl acetate and allowed to sit overnight with occasional mixing. Ethyl acetate fractions were collected using a separatory funnel and dried under vacuum in a Rotavapor® rotary evaporator (Buchi AG, Switzerland) at 32°C. Metabolites were resuspended in DMSO and filtered through a 0.45 μm Durapore™ filter (Millipore, USA). Extracts were reassessed for antibiofilm activity prior to LC-MS/MS analysis. Briefly, 100µL solubilized ethyl acetate extracts were mixed with 15µL of 2% acetonitrile and 0.1% formic acid prior to 10µL injections on a 25cm × 75µm column. MS/MS data was collected on charge states 1-6 using an Orbitrap analyzer.

### Fractionation of Ethyl Acetate Organic and 100%P1 Extracts

To simplify the ethyl acetate extract, flash chromatography was performed on a Teledyne Isco CombiFlash NextGen 100, fitted with a C18 column. The extract was dissolved in 25% methanol and injected onto the column. A stepwise gradient of water (A) and methanol (B) from 15%B, 25%B, 50%B, 75%B, to 100%B was used to separate the extract. Fractions were collected using the automatic fraction collector. During the stepwise gradient, individual peaks were collected separately from the rest of the flow through to provide a fraction enriched with those compounds. These peaks were labeled as X%P1 for the first peak separated in any given fraction. Divided fractions were combined and dried in vacuo using a rotary evaporator. To further separate 100%P1, this fraction (221 mg) was separated on a reversed-phase open column (RediSep Rf C18 50g; 60 to 100 % aq. methanol) to generate two subfractions containing only surfactins (100%P1-A) or only the unique compound (100%P1-B). Fraction 100%P1-B (142 mg) was purified using a Dionex UltiMate 3000 HPLC system, using a Luna C18 column (250 mm × 15 mm, Phenomenex), running acetonitrile with 0.1% formic acid and H2O with 0.1% formic acid as the mobile phase.

### LC-MS/MS Analysis

Fractions and isolated peaks were dried in vacuo, resuspended in 50% methanol, and analyzed on a Q-Exactive quadrupole orbitrap mass spectrometer (Thermo Scientific, USA) coupled to a UPLC (Dionex). The UPLC method was 5% methanol for 0.5 minutes followed by a linear gradient from 5% methanol to 97% methanol over 16 minutes. The 97% methanol wash was held for two minutes before switching back to 5% methanol over 0.5 minutes and re-equilibrating at 5% methanol for 1 minute. The UPLC flow rate was 0.35 mL/min over a Phenomenex Kinetex XB-C18 chromatography column with dimensions 2.1 x 100 mm and particle size 2.6 micron. The mass spectrometer scanned from 200 to 2000 m/z in positive mode and ion fragmentation was achieved using stepped normalized collision energy of 30, 35, and 40%. The data was collected in profile mode and manually inspected and filtered using MzMINE2. MzMINE2 was used to create an aligned feature list and quantification table of the mass spectral data of the biologically active peak. These files were used to perform feature-based molecular networking through GNPS^61^. The GNPS spectral libraries were searched for matches to submitted MS/MS spectra with a cosine score threshold of 0.7 and minimum of 6 matched peaks to the library spectra. The resulting network file was visualized and analyzed using Cytoscape v3.8.0.

### Cytokine and cytotoxicity assays

Gibco™ Dulbecco’s Modified Eagle Medium (DMEM) (Thermo Scientific, USA) consisting of 10% fetal bovine serum and 0.1% Gibco™ Antibiotic-Antimycotic (10,000 units/mL penicillin, 10,000ug/mL streptomycin, 25ug/mL amphotericin B) (Thermo Scientific, USA). Cells were cultured for 7 days at 37°C in a carbogen (95% O2, 5% CO2) atmosphere, with the culture medium being changed every two to three days until cells were confluent. For CCL81 cell lines, approximately 2 × 10^4^ cells were seeded into new 96 well plates (Corning, USA) and allowed to form monolayers for 48 hours prior to cytotoxicity testing. For HT29 cells, approximately 2 × 10^4^ cells were seeded into new 24-well plates (Corning, USA) and allowed to differentiate for 33 days prior to treatment. After removing media, cells were washed twice with Gibco™ PBS, pH 7.4 (Thermo Scientific, USA) and equilibrated as previously described before adding bacteria^62,63^. Vero cells were equilibrated in EC buffer (135mM NaCl, 15mM HEPES, 1mM MgCl2, 1mM CaCl2) for four hours before adding bacteria. HT29 cells were equilibrated in DMEM without antibiotic for four hours prior to adding bacteria. Exponential phase *S. aureus* ATCC 29213 cultures grown in 1.5% TSB + 0.3% glucose media were standardized to OD_600_ 1.0, centrifuged, washed with DPBS, and applied to appropriate test wells at a final concentration of ∼6 x 10^7^ CFU/well. After treatment, cells were centrifuged at 600g for 5 minutes. Cytotoxicity was determined using a lactate dehydrogenase (LDH) assay kit according to the manufacturer’s protocol (Abcam, UK). Using HT29 cells, the human pro-inflammatory TNF-α, the dual functioning IL-6, and the anti-inflammatory IL-10 ELISAs were performed using Quantikine® Colorimetric Sandwich ELISA Kits (R&D Systems, USA) according to the manufacturer’s protocol. All treatments were performed in triplicate and assayed in duplicate.

### Statistical analysis

Statistical analyses were performed using GraphPad Prism 9.5.0 (San Diego, CA). Differences where p<0.05 were considered significant. Differences in biofilm inhibition were analyzed using a 2-tailed unpaired t-test. Differential gene expression was analyzed using one-way ANOVA, followed by Tukey’s multiple comparisons using log2 transformed data. Cytokine levels were compared between the differently treated cell lines using Welch’s non-parametric tests.

### Data Availability

Genome assemblies were deposited in NCBI under BioProject accession number PRJNA988465. All LC-MS/MS data were deposited in the Center for Computational Mass Spectrometry’s MassIVE database under project number MSV000092675.

## Results

### B. subtilis 6D1 exhibits antibiofilm activity against S. aureus and harbors unique genetic traits not observed in closely related B. subtilis strains

High-throughput screening of 1123 presumptive *Bacillus* strains revealed *B. subtilis* 6D1 exhibited the strongest antibiofilm activity among the sixteen most potent environmentally sourced strains (Fig. 1A). The genomic diversity among *B. subtilis* strains can be leveraged to understand how certain strain elements contribute to their antimicrobial production potential^27^. Therefore, whole genome sequencing, phylogenomic, and pangenome analyses of *B. subtilis* 6D1 were performed to search for unique genomic traits that might explain this strain’s prominent antibiofilm activity. Genomic assembly yielded two contigs – the 4MB chromosome and an 84kB plasmid similar to pBS32, a large ancestral plasmid rarely found in environmental isolates^64,65^ (Fig. 1B). The entire genome contained 4,156,213 base pairs with 43.65% GC content. A total of 4,309 protein coding genes (CDS), 88 transfer RNA (tRNA) genes, and 30 ribosomal RNA (rRNA) genes were predicted. Of the 4,309 CDS predicted, 708 were hypothetical proteins, and 3,601 proteins had functional assignments – these included 1,055 proteins with Enzyme Commission (EC) numbers, 881 with Gene Ontology (GO) assignments, and 776 proteins that were mapped to KEGG pathways. Sixteen antimicrobial resistance genes were identified only in the chromosome and were not flanked by any predicted mobile genetic elements (Fig. 1B).

Taxonomic identification and phylogenetic analysis confirmed this genome was a member of the *B. subtilis* group and was most similar to *B. subtilis* 75, and *B. subtilis* PTA-271, strains sourced from the rhizosphere of high performing plant species capable of reducing plant pathogen colonization (Fig. S2). Phylogenomic analysis revealed *B. subtilis* 6D1 shares >0.95% identity with a recently evolved group of *B. subtilis* genomes (Fig. 1C); however, this strain encodes an additional 156 singleton genes that were not present in these closely related strains (Fig. 1D, Table S3). Many of the singleton genes were of unknown function, but genes encoding a glycosyl hydrolase were identified. These enzymes hydrolyze the glycosidic bonds between sugars, such as those found within the exopolysaccharides of biofilm matrices and have been shown to reduce biofilm biomass by weakening the matrix and inducing bacterial dispersal^66,67^. Interestingly, glycosyl hydrolases can inhibit and degrade Staphylococcal and Pseudomonal biofilms and improve clearance of these pathogens when applied in conjunction with antibiotics^67^. The second largest group of singleton genes identified were predicted to be of viral origin, including prophage genes from the family Siphoviridae. Genome mapping further confirmed the presence and location of a 25kB and 27kB prophage region predicted to belong to the Siphoviridae and Myoviridae family, respectively (Fig. 1B). The *B. subtilis* 6D1 strain appeared to maintain many ancestral *B. subtilis* genes (Fig. 1D), including a PBS32-like plasmid (Fig. 1B); however, our results suggest this strain may have recently acquired foreign genetic elements from phage that increased their ability to degrade pathogenic biofilms.

### B. subtilis 6D1 inhibits S. aureus biofilm growth but not planktonic growth

To further explore the mechanisms by which *B. subtilis* 6D1 inhibits *S. aureus* biofilms, cell-free extracts (CFEs) were applied to clinical methicillin susceptible and methicillin resistant *S. aureus* strains and evaluated for antimicrobial and antibiofilm activity. Compared to 0.5X MIC sulfamethoxazole/trimethoprim, *B. subtilis* 6D1 CFE applied at 10% v/v significantly reduced biofilm formation of *S. aureus* ATCC 29213 and both clinical strains (Fig. 2B) without inhibiting planktonic growth (Fig. 2A). Confocal image analysis confirmed antibiofilm activity of 10% v/v *B. subtilis* 6D1 CFE, reducing mean Syto9 intensity 77% compared to untreated wells (Fig 2D-E). Macroscopic observation of *S. aureus* ATCC 29213 biofilms grown for 24 hours and washed prior to SYTO-9 staining corroborated microscopic imaging (Fig. 2F). Subsequent size fractionation analyses revealed antibiofilm activity was maintained in CFE fractions <3kDa (Fig. S1), suggesting small peptides, rather than glycosyl hydrolases, are likely driving this effect. Competition experiments in planktonic and biofilm environments also revealed *B. subtilis* 6D1 outcompetes *S. aureus* ATCC 29213 in a biofilm but not in a planktonic environment (Fig. 2C). These data suggest *B. subtilis* 6D1 inhibits *S. aureus* biofilm formation during cell-to-cell interactions, confirming that antibiofilm activity is not simply an artifact of monoculture CFE preparations. Taken together these results demonstrate compounds produced by *B. subtilis* 6D1 are capable of inhibiting biofilm formation in *S. aureus*.

**Figure 2.**
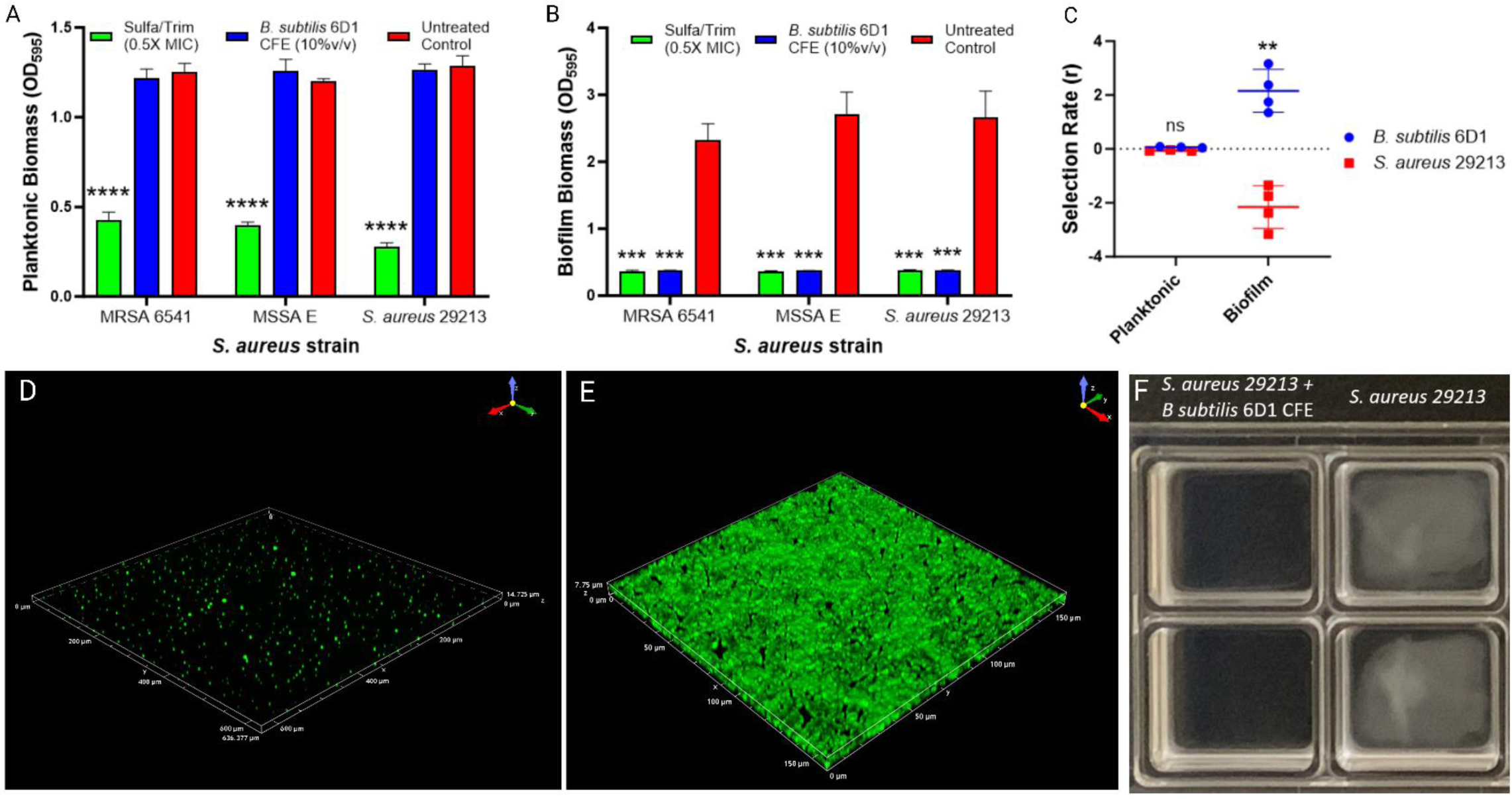
*B. subtilis* 6D1 inhibits *S. aureus* biofilm but not planktonic growth. **(A-B)** Compared to 0.5X MIC of Sulfamethoxazole/Trimethoprim, addition of 10% v/v *B. subtilis* 6D1 cell-free extract inhibits methicillin susceptible (MSSA) and methicillin resistant (MRSA) *S. aureus* biofilm growth, but not planktonic growth. (***) P<0.001, (****) P<0.0001 compared to the untreated control. **(C)** Competition experiments performed in planktonic and biofilm conditions for 24 hours show *B. subtilis* 6D1 outcompetes *S. aureus* in a biofilm but not in a planktonic environment, (**) P<0.01. **(D)** SYTO-9 staining visualized by confocal microscopy confirmed antibiofilm activity of 10% v/v *B. subtilis* 6D1 CFE **(E)** compared to an untreated control of *S. aureus* ATCC 29213. **(F)** Macroscopic observation of *S. aureus* ATCC 29213 biofilms grown for 24 hours and washed prior to Syto9 staining.

### B. subtilis 6D1 cell-free extracts reduce S. aureus biofilm growth, disassemble mature biofilm, and improve biofilm inhibition when applied in conjunction with low doses of antibiotics

To assess which stages of the *S. aureus* ATCC 29213 biofilm growth cycle^68^ were impacted by *B. subtilis* 6D1, 10% v/v CFE was applied at various timepoints throughout biofilm growth. If applied within the first six hours, the CFE reduced biofilm growth (Fig. 3B), and biofilm formation did not recover after initial CFE exposure (Fig. 3A). These results suggest *B. subtilis* 6D1 produces soluble elements that inhibit biofilm adherence and early maturation – processes typically under the control of the Agr QS system^69^. Furthermore, addition of *B. subtilis* 6D1 CFE reduced *S. aureus* biofilm growth (Fig. 3C) and disrupted mature biofilm (Fig. 3D) in a concentration dependent manner. These data indicate a positive correlation between active compound abundance and antibiofilm activity.

**Figure 3.**
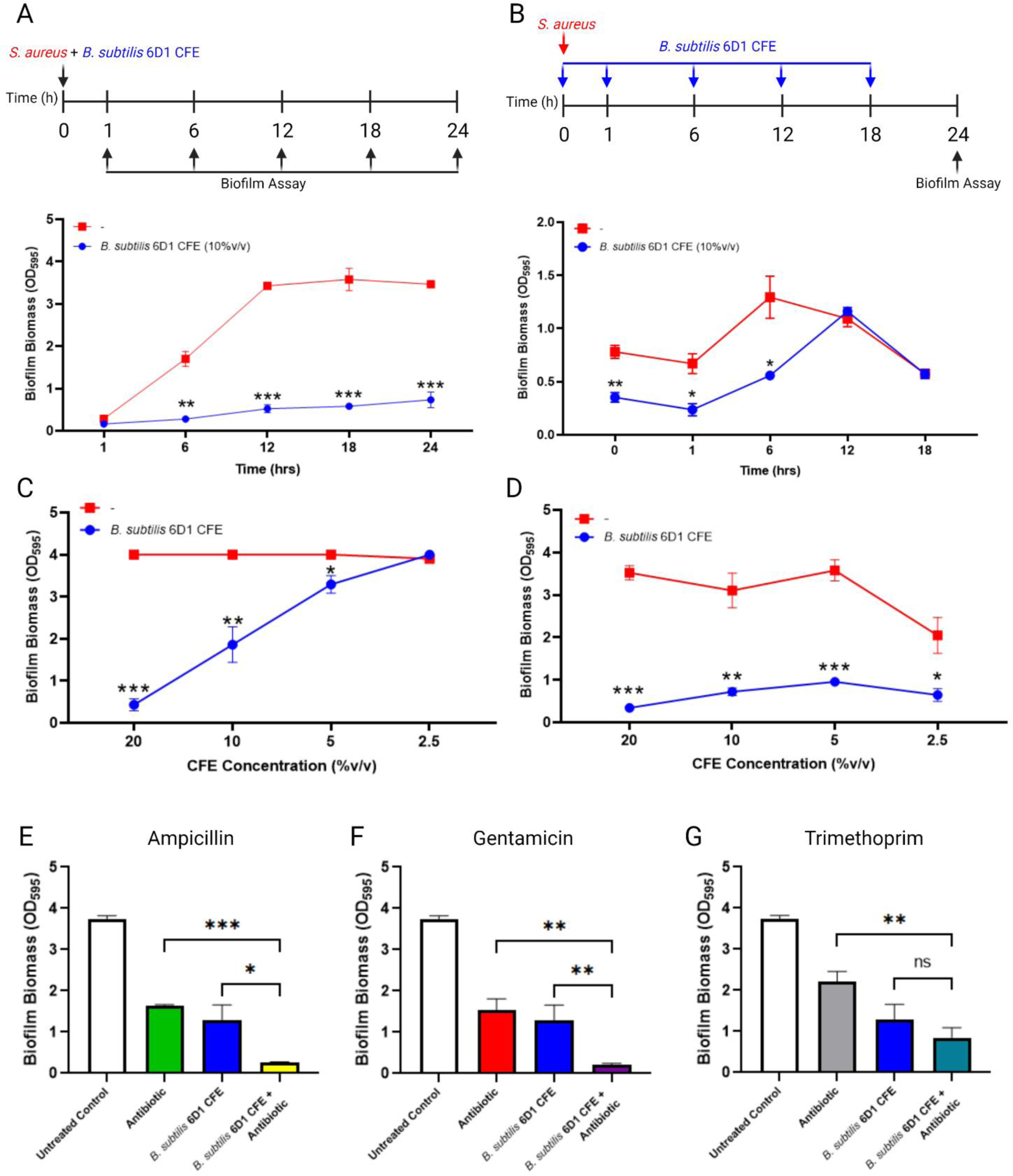
*Bacillus subtilis* 6D1 cell-free extracts (CFEs) reduce *S. aureus* ATCC 29213 biofilm growth, disassemble mature biofilm, and improve biofilm inhibition when applied in conjunction with low doses of antibiotics. **(A)** *S. aureus* (5×10^5^ CFU/ml) was grown on polystyrene plates at 37°C for 1, 6, 12, 18, or 24 hours in the presence or absence of *B. subtilis* 6D1 CFE (10% v/v) **(B)** *S. aureus* (5×10^5^ CFU/mL) was grown on polystyrene plates at 37°C for 0, 1, 6, 12 or 18 h, followed by treatment with *B. subtilis* 6D1 CFE (10% v/v) at 37°C for up to 24h **(C)** *B. subtilis* 6D1 CFE (20%, 10%, 5%, or 2.5% v/v) was applied concurrently with *S. aureus* at T0 prior to staining biofilm at 24h **(D)** *B. subtilis* 6D1 CFE (20%, 10%, 5%, or 2.5% v/v) was applied to *S. aureus* 24h biofilms and incubated at 37°C shaking at 100rpm for 1 hour **(E-G)** *B. subtilis* 6D1 CFE (10%v/v) applied in conjunction with 0.5μg/mL antibiotic at 0h prior to measuring *S. aureus* biofilm growth at 24h; (*) P<0.05; (**) P<0.01; (***) P<0.001

Next, we tested whether application of *B. subtilis* 6D1 CFE improves antibiotic efficacy against biofilm growth if applied in conjunction with low doses of antibiotics. Ampicillin, gentamicin, and trimethoprim were selected for both their clinical relevance in treating Staphylococcal infections and their different antimicrobial mechanisms– ampicillin interferes with cell wall synthesis, gentamicin inhibits protein translation, and trimethoprim prevents DNA synthesis. To quantify synergistic, additive, indifferent, or antagonistic effects brought on by the addition of *B. subtilis* 6D1 CFE, minimum biofilm inhibition concentrations (MBIC) and minimum inhibitory concentrations (MIC) were determined for each antibiotic to identify the lowest permissible concentration that would allow *S. aureus* ATCC 29213 planktonic and biofilm growth. Application of 10% v/v *B. subtilis* 6D1 CFE in conjunction with ampicillin (FICI = 0.34 ± 0.08) and gentamicin (FICI = 0.28 ± 0.04) synergistically reduced *S. aureus* ATCC 29213 biofilm formation and possessed an additive inhibitory effect when applied in conjunction with trimethoprim (FICI = 0.87 ± 0.08) (Fig. 3E-G).

### B. subtilis 6D1 modulates gene expression associated with S. aureus quorum sensing

*S. aureus* quorum sensing involves the recognition of AIPs by their cognate AgrC receptors which prompts the phosphorylation of AgrA and the subsequent expression of the Agr operon^20^. RT-qPCR experiments were conducted to investigate how *B. subtilis* 6D1 or its associated cell-free extract altered the expression of *S. aureus* biofilm and QS related genes (Table S2). Our data demonstrated that both 10% v/v CFE and *B. subtilis* 6D1 cells applied in a 1:1 ratio with *S. aureus* ATCC 29213 increased expression of *agrA*, *RNAIII*, and *hld* – all genes under the control of the Agr QS system (Fig. 4). Both *B. subtilis* 6D1 and the CFE increased *agrA, RNAIII,* and *hld* expression ranging from 4-fold to 29-fold. *AgrA*, *RNAIII*, and *hld* are all located in the *S. aureus* Agr operon and when upregulated, transition cells from a biofilm lifestyle to a planktonic lifestyle^69^. Other genes also upregulated in response to *B. subtilis* 6D1 exposure included the global stress response regulator, *sigB*, and *saeR*, the response regulator of a two component signal transduction system that regulates the expression of numerous virulence factors in *S. aureus*, including surface-bound and secreted proteins^70^. Compared to the 1:1 *B. subtilis* 6D1 cell-to-cell treatment, *B. subtilis* 6D1 CFE elicited a 9.2-fold and 6.3-fold stronger expression response for both *sigB* and *saeR*, respectively.

**Figure 4.**
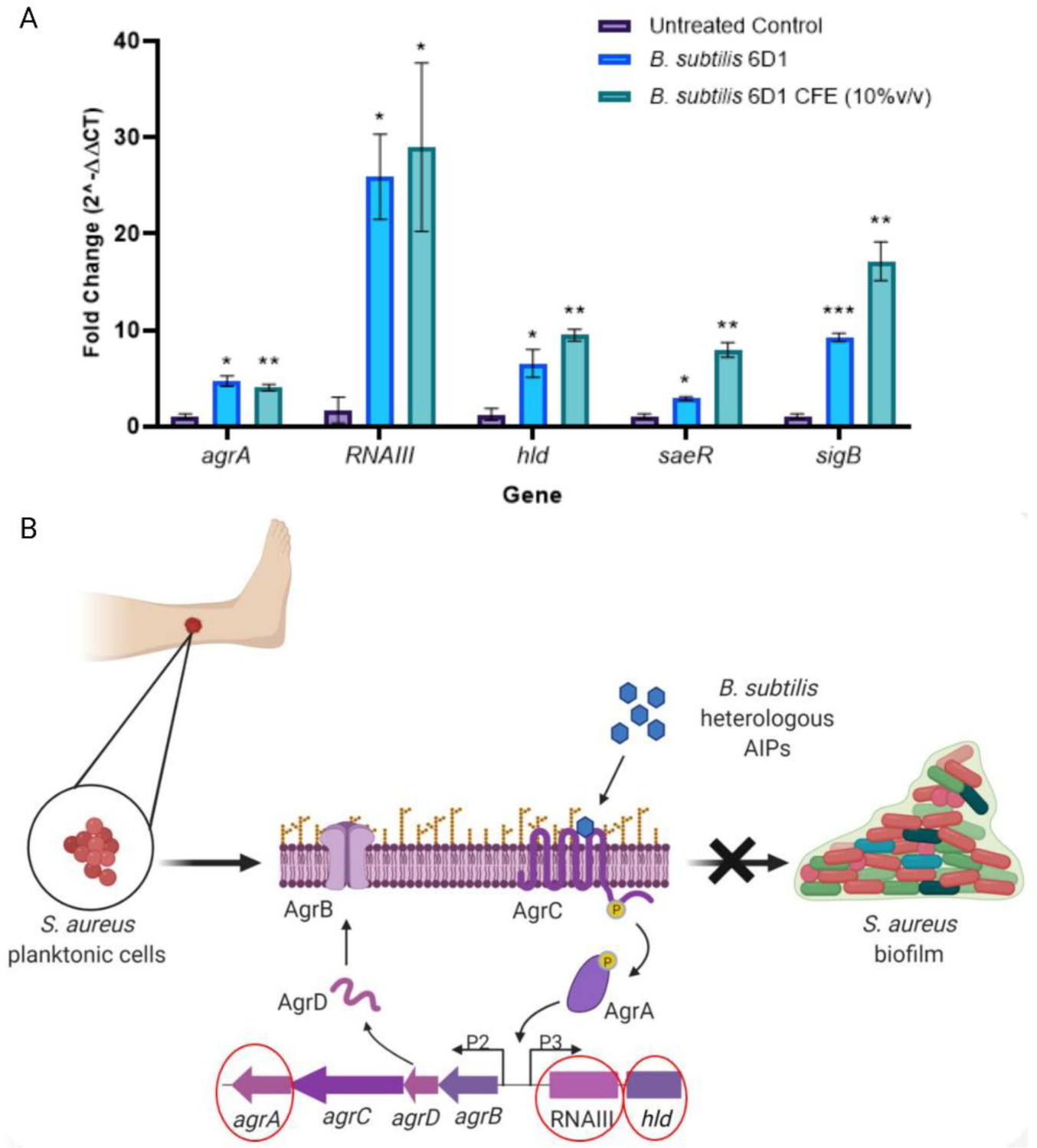
*B. subtilis* 6D1 modulates gene expression associated with *S. aureus* quorum sensing. **(A)** *B. subtilis* 6D1 (blue) and 10% v/v *B. subtilis* 6D1 cell-free extract (teal) were applied to *S. aureus* ATCC 29213 for 24 hours before RNA extraction and expression analysis via RT-qPCR **(B)** Proposed model of *S. aureus* QS interference by *B. subtilis* 6D1*. B. subtilis* derived AIPs bind to AgrC and trigger phosphorylation and activation of DNA binding protein AgrA, (*) P<0.05; (**) P<0.01; (***) P<0.001 compared to the untreated control.

To explore which CFE compounds might be driving this activity, the *B. subtilis* 6D1 genome was mined for biosynthetic gene clusters (BGCs) that might produce secondary metabolites. Seven BGCs were identified and subsequently mapped to reference BGCs identified in the undomesticated *B. subtilis* NCIB 3610 strain. Of these seven predicted BGCs, five were found to possess 100% similarity with reference clusters: bacilysin, bacillaene, bacillibactin, subtilosin A, and sporulation killing factor (Fig. S3). Interestingly, two BGCs showed 78% similarity to surfactin and 93% similarity to fengycin, non-ribosomal peptides with previously demonstrated quorum sensing interference and antibiofilm capabilities^71,72^. However, subsequent biofilm inhibition experiments using commercial fengycin obtained from *B. subtilis* revealed this compound increased biofilm formation in multiple Agr backgrounds (Fig. S4). Additional BGC mapping determined that the initially identified fengycin cluster was instead, plipistatin, a very similar lipopeptide but with a slightly different structural moiety^73^. Furthermore, compared to the plipistatin reference BGC, the *B. subtilis* 6D1 BGC has lost both *ppsA* and *ppsB* non-ribosomal peptide synthetase (NRPS) genes (Fig. S3), suggesting either this metabolite is structurally distinct from plipistatin or this BGC is transcriptionally inactive^74^. Compared to the surfactin reference BGC, *srfAA* and *srfAB* NRPS genes identified in *B. subtilis* 6D1 shared 75% and 77% identity, respectively (Fig. S3). *B. subtilis* 6D1 also harbors an additional ABC transporter downstream of *yciC* not identified in the reference BGC. Taken together, we conclude that *B. subtilis* 6D1 harbors biosynthetic gene clusters that closely resemble surfactin BGCs – a peptide capable of inhibiting *S. aureus* biofilm and interfering with Agr quorum sensing. Therefore, liquid-liquid extractions were performed to determine if these lipopeptides are produced, and whether they contribute towards the antibiofilm activity observed.

### Peptides isolated from B. subtilis 6D1 cell-free extracts inhibit biofilm growth in all Agr backgrounds

**(A)** *S. aureus* strains can harbor one of four distinct Agr QS systems, and though each Agr system has been associated with a different disease etiology, the Agr QS system in each group is responsible for regulating virulence factor gene expression and facilitating biofilm formation and dispersal^14,75^. To assess whether *B. subtilis* 6D1 lipopeptides were driving *S. aureus* antibiofilm activity, a crude ethyl acetate extract resuspended in DMSO was assessed for its antibiofilm activity against *S. aureus* strains possessing different Agr backgrounds. This extract exhibited dose-dependent activity, and its ability to inhibit biofilm formation in an Agr null background suggests that it may be *Bacillus* lipopeptides, not self-stimulated production of AgrD, driving the expression of genes within the *agr* operon (Fig. 5A).

**Figure 5.**
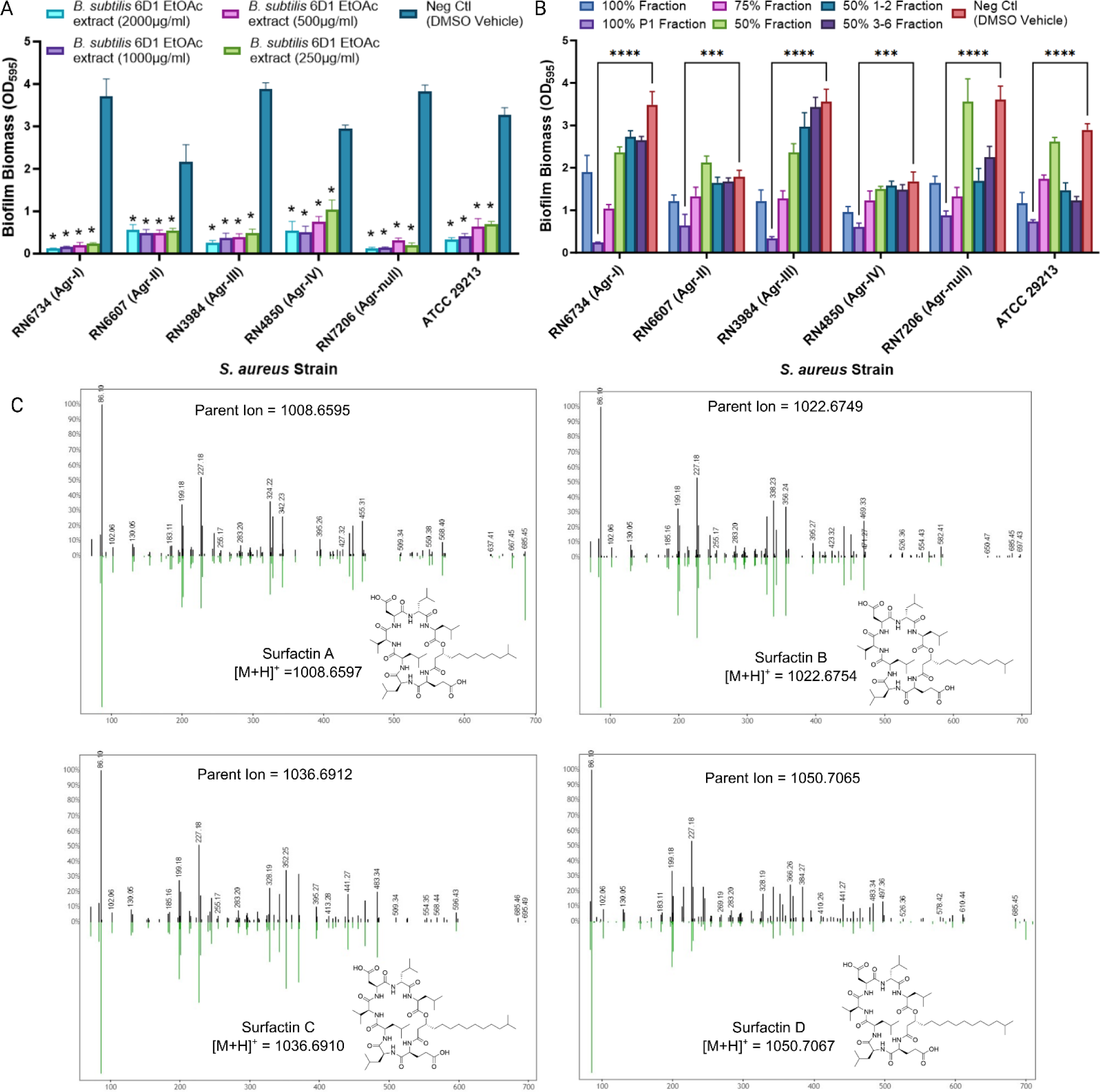
*B. subtilis* 6D1 derived peptides inhibit biofilm growth in all Agr backgrounds. **(A)** *B. subtilis* 6D1 ethyl acetate extracts exhibited dose dependent antibiofilm activity against *S. aureus* strains with different AgrC receptors (*agr* I-IV) and a strain incapable of producing AgrD (RN7206) (*) P<0.0001 **(B)** Antibiofilm activity of *B. subtilis* 6D1 ethyl acetate extracts (500μg/ml) further separated by combiflash fractionation. (***) P<0.001, (****) P<0.0001 **(C)** LC-MS/MS spectra of *B. subtilis* 6D1 fraction 100%P1 m/z values that matched known molecules in the GNPS database. Black spectra represent peaks identified in fraction 100% P1 and green spectra represent GNPS database spectra. The x-axis represents m/z and the y-axis represents the relative abundance of the various ions.

*B. subtilis* 6D1 ethyl acetate extracts were separated further by flash chromatography using a stepwise gradient of water and methanol to obtain six distinct fractions with unique LC-MS/MS spectra. After drying, concentrated fractions were dissolved in DMSO and again evaluated for their ability to inhibit *S. aureus* biofilm formation. Compared to DMSO vehicle controls, fraction 100%P1 harbored the strongest antibiofilm activity among the six fractions (Fig. 5B). Though other fractions harbored antibiofilm activity, namely 100% and 75% fractions, 100%P1 yielded the most consistent antibiofilm activity across all Agr backgrounds. Feature based molecular networking through GNPS confirmed the presence of surfactin A-D in fraction 100%P1 (Fig. 5C). A smaller sub-network with no known matches to GNPS database spectra was also identified in this fraction, and subsequent separation of this compound from the surfactins revealed it, too, possessed potent antibiofilm activity (Fig. S5).

Both *B. subtilis* NCIB 3610 and another environmental *B. subtilis* strain, 9B5, harboring moderate antibiofilm activity were analyzed via LC-MS/MS alongside *B. subtilis* 6D1, but neither strain possessed this unknown sub-network. Collectively, we conclude that lipopeptides produced by *B. subtilis* 6D1, namely surfactin A-D and a novel compound, inhibit biofilm growth in all four *S. aureus* Agr backgrounds and in an Agr null strain incapable of producing its own autoinducing peptide, AgrD.

### B. subtilis 6D1 100%P1 fraction exhibits stronger antibiofilm activity compared to HPLC grade surfactin obtained from B. subtilis

A series of biofilm inhibition experiments were performed comparing the 100%P1 fraction and commercial HPLC grade surfactin produced by *B. subtilis* (Sigma-Aldrich, USA). Both 100%P1 and commercially obtained surfactin were standardized in DMSO to 10mg/mL working stock concentrations to ensure that both identical DMSO volumes and metabolite concentrations were applied to test conditions. Additional DMSO titrations confirmed volumes used to deliver these peptides did not increase cell death compared to the untreated biofilm controls (Fig. S6). Though both commercial surfactin and fraction 100%P1 successfully inhibited *S. aureus* biofilm formation, preliminary titration experiments revealed that at 500μg/mL, 100%P1 harbored stronger antibiofilm activity (94 ± 0.7%) than commercial surfactin alone (73 ± 2.6%) (Fig. 6A). Confocal image analysis confirmed these findings – 100%P1 treated biofilms reduced SYTO9 mean fluorescence intensity 72% more than commercial surfactin (Fig. 6B). Based on these observations, a 500μg/mL concentration was used in all ensuing experiments. At this concentration, *B. subtilis* 6D1 fraction 100%P1 possessed stronger antibiofilm activity against all Agr background strains with biofilm percent inhibition ranging from 88-97% while biofilm inhibition ranged from 74-92% for surfactin alone (Fig. 6C). This trend persisted when evaluating clinical *S. aureus* isolates. *B. subtilis* 6D1 fraction 100%P1 exhibited biofilm inhibition ranging from 82-93% while surfactin biofilm inhibition was much less consistent, ranging from 46-85% in the same isolates (Fig. 6D). Further separation of fraction 100%P1 revealed that both the surfactin mixture and the unique compound found in this fraction exhibited strong biofilm inhibition and disruption activity (Fig. S5). Therefore, it is possible this collection of 100%P1 compounds are either additively or synergistically increasing *S. aureus* antibiofilm activity compared to commercial surfactin.

**Figure 6.**
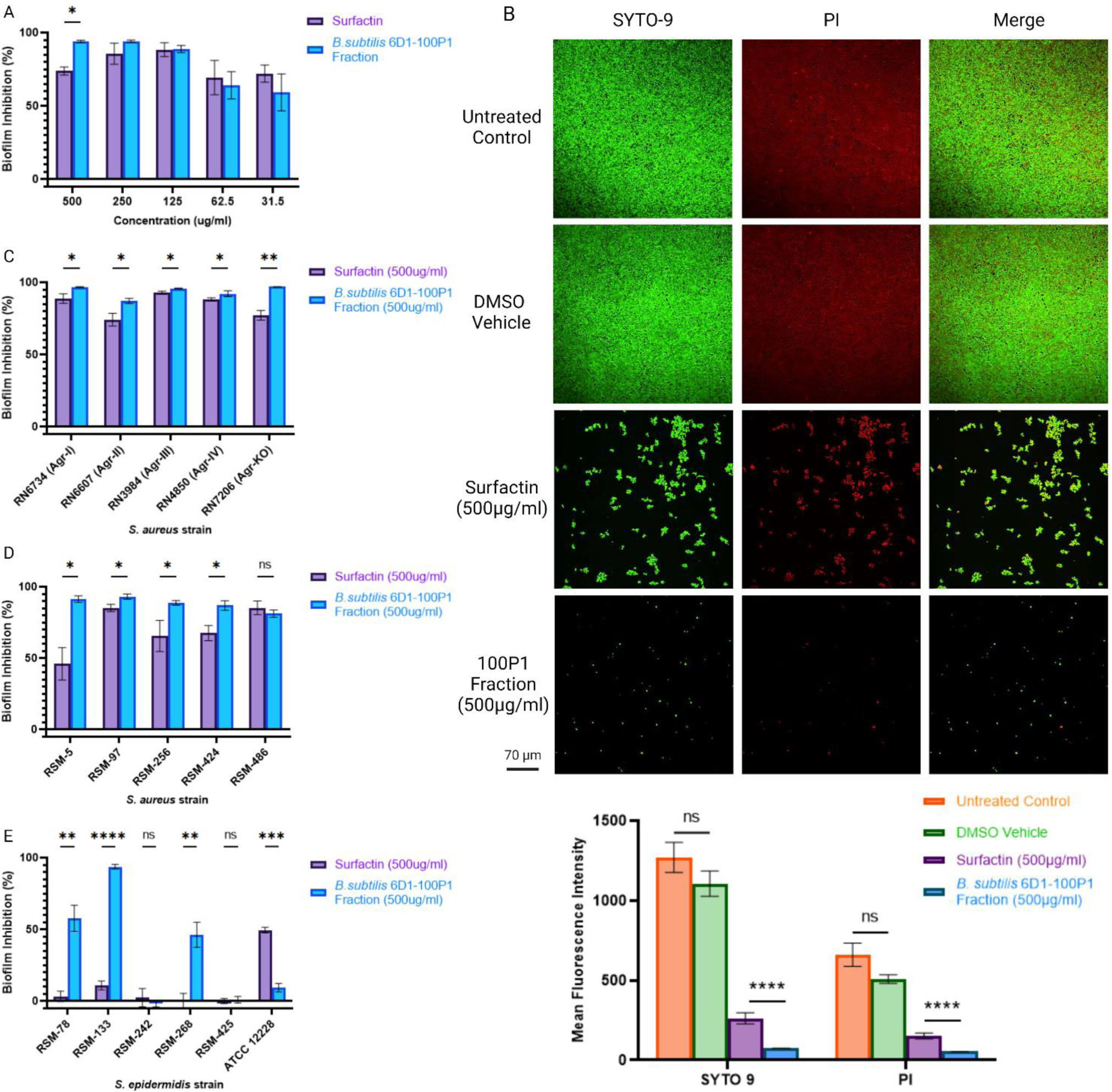
*B. subtilis* 6D1 100P1 fraction exhibits unique antibiofilm activity compared to commercial HPLC grade surfactin obtained from *B. subtilis*. **(A)** Antibiofilm activity of titrated concentrations of commercial surfactin and *B. subtilis* 6D1 fraction 100% P1 against *S. aureus* ATCC 29213. **(B)** Live (SYTO9)/dead(PI) staining and confocal image analysis comparing the arithmetic mean fluorescence intensity of *S. aureus* ATCC 29213 biofilms treated with 500μg/mL commercial surfactin and *B. subtilis* 6D1 fraction 100% P1, scale bar = 70μm. **(C)** Antibiofilm activity of commercial surfactin and *B. subtilis* 6D1 fraction 100% P1 against *S. aureus* strains with different AgrC receptors (*agr* I-IV) and a strain incapable of producing AgrD (RN7206) **(D)** Antibiofilm activity of commercial surfactin and *B. subtilis* 6D1 fraction 100% P1 against clinical *S. aureus* (E) and *S. epidermidis* strains. (*) P<0.05; (**) P<0.01; (***) P<0.001; (****) P<0.0001

Interestingly, differences between 100%P1 and surfactin were more apparent when evaluating biofilm inhibition activity against coagulase negative *Staphylococcus epidermidis*, another species with a functioning Agr QS system that can form biofilm on indwelling medical devices^76,77^. Fraction 100%P1 successfully inhibited 3/5 clinical *S. epidermidis* isolates, however biofilm inhibition did not exceed 10% for any of the five clinical strains treated with commercial surfactin (Fig. 6E). Conversely, surfactin only inhibited biofilm formation of *S. epidermidis* ATCC 12228, while fraction 100%P1 did not have this effect. Based on this data, we conclude that at 500μg/mL concentrations, *B. subtilis* 6D1 fraction 100%P1 comprised of multiple surfactin isoforms and a unique compound exhibits stronger antibiofilm activity than commercial surfactin against *S. aureus* strains.

### B. subtilis 6D1 reduces S. aureus ATCC 29213 virulence in a human intestinal cell line

It is known that staphylococcal virulence factor production and host cell invasion is mediated by Agr-dependent processes^78^, so it is important to establish whether compounds altering *agr* expression also modify *S. aureus* virulence^79^. Therefore, both Vero (CCL81) and HT29 cell lines were used to assess the cytotoxic and ameliorative effects of *B. subtilis* 6D1 CFE, fraction 100%P1, and commercial surfactin in the presence and absence of an *S. aureus* challenge. Both 100%P1 and commercially obtained surfactin were standardized in DMSO to 10mg/mL working stock concentrations to ensure that both identical DMSO volumes and fraction dry weight/volume concentrations were applied to test conditions. At 250μg/mL, fraction 100%P1 reduced *S. aureus* induced cytotoxicity from 43.4 ± 12.82% to 31.8 ± 2.24%, however, this protective effect was lost at concentrations below 200μg/mL. In the absence of a *S. aureus* challenge, fraction 100%P1 was less cytotoxic than commercial surfactin and exhibited a decreasing trend in line with the DMSO vehicle (Fig. 7C). In fact, fraction 100%P1 was significantly less cytotoxic than commercial surfactin at all concentrations except 200μg/mL (P=0.09). Interestingly, reduced DMSO concentrations also appeared to protect Vero cells, significantly inhibiting *S. aureus-*induced cytotoxicity by 24% at 50μg/mL (Fig. 7B). This phenomenon has been observed in other cell culture studies, where low concentrations of DMSO elicit an immunomodulatory response and increased cellular activation, however these results are not universal and appear to be cell line specific^80,81^.

**Figure 7.**
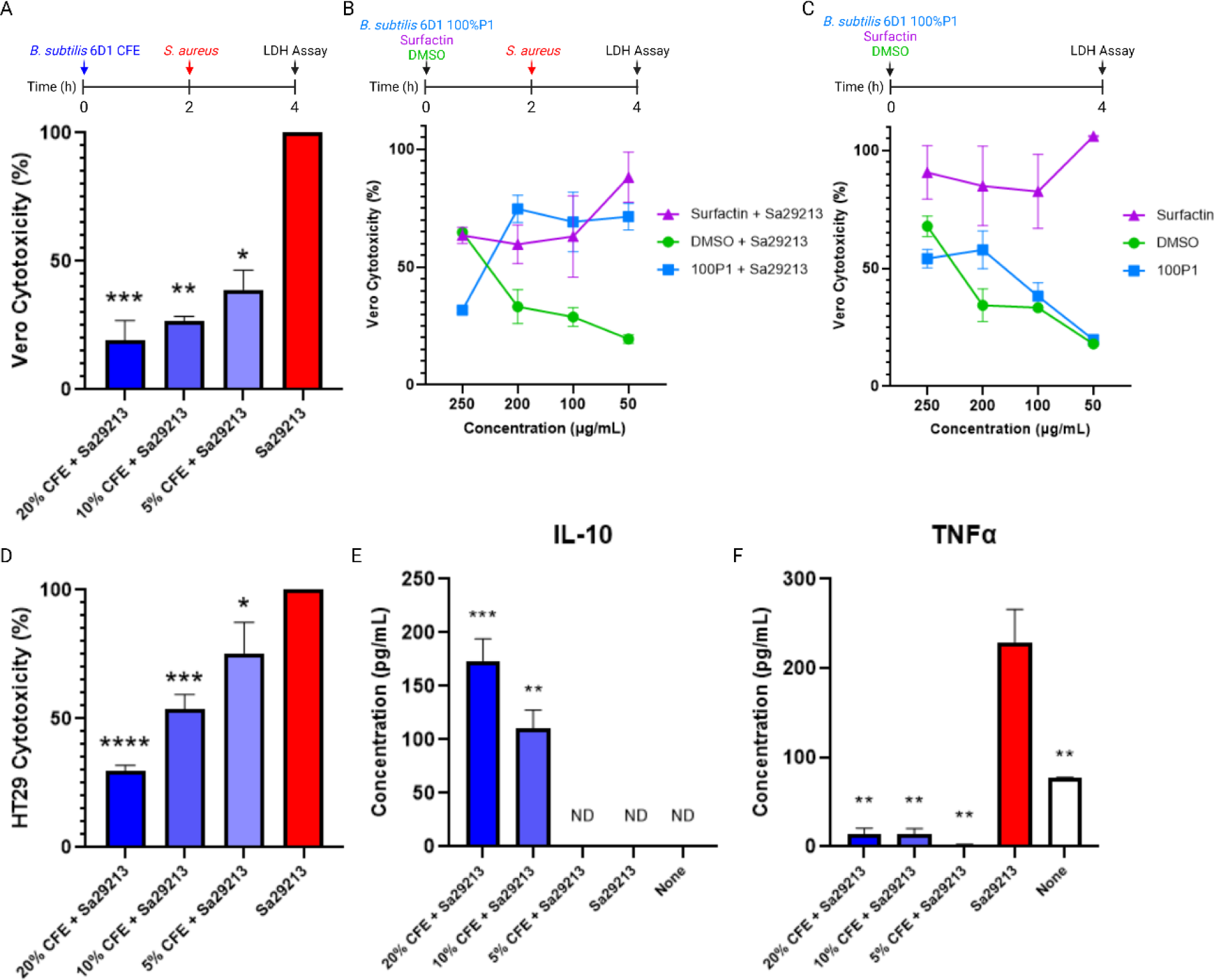
*B. subtilis* 6D1 reduces *S. aureus* ATCC 29213 virulence in a human intestinal cell line. **(A)** Vero cells pretreated for two hours with increasing concentrations of *B. subtilis* 6D1 CFE reduce cytotoxicity of *S. aureus* ATCC 29213 (Sa29213) induced cell death; data normalized to Sa29213 positive infection control. **(B)** Cytotoxicity of Vero cells pretreated for two hours with increasing concentrations of fraction 100%P1 and surfactin delivered in DMSO in the presence **(C)** and absence of a Sa29213 challenge; data normalized to LDH positive control. **(D)** HT29 cells pretreated for two hours with increasing concentrations of *B. subtilis* 6D1 CFE inhibit Sa29213 induced cytotoxicity; data normalized to Sa29213 positive infection control **(E)** HT29 cells pretreated for two hours with *B. subtilis* 6D1 CFE increase production of the anti-inflammatory cytokine IL-10 **(F)** and inhibits the production of the pro-inflammatory cytokine TNFα brought on by Sa29213 challenge; (*) P<0.05; (**) P<0.01; (***) P<0.001 (****) P<0.0001, ND = not detected

Next, we investigated whether *B. subtilis* 6D1 CFE possessed any additional protective effects beyond those observed using the refined 100%P1 fraction. Surprisingly, *B. subtilis* 6D1 CFE inhibited *S. aureus* induced Vero cell toxicity in a dose-dependent manner (Fig. 7A). Based on these data we sought to investigate whether the ameliorative effects observed with *B. subtilis* 6D1 CFE in Vero cells were maintained in a human intestinal cell line (HT29) and quantify specific immunological markers that may be driving this effect. Application of CFE exhibited a similar dose dependent protective effect in HT29 cells challenged with *S. aureus* 29213 (Fig. 7D). Further investigation revealed this protective effect was assisted by the production of anti-inflammatory cytokine IL-10 and reduced production of the pro-inflammatory cytokine, TNFα (Fig. 7E-F). Together, these data demonstrate that while fraction 100%P1 exerts strong antibiofilm activity, the mixture of compounds in *B. subtilis* 6D1 CFE may possess broader beneficial properties to help combat a variety of *S. aureus* virulence strategies.

## Discussion

As a first step in determining novel, *B. subtilis*-derived products capable of reducing virulence and improving antibiotic efficacy against *S. aureus*, it is essential to source probiotic strains from environments where frequent interactions with *S. aureus* might occur^82^. In our case, 1123 *Bacillus* strains were sourced from bovine milk and feces and a variety of environments impacted by multiple dairy operations comprising nearly 100,000 dairy cows in Kewaunee county, WI^83^. We hypothesized that *Bacillus* strains obtained from these environments had likely evolved coercion tactics to compete with bovine *S. aureus* strains through inhibition of biofilm formation^84^. In line with this hypothesis, we isolated *B. subtilis* 6D1 and determined this strain possessed 156 unique genes not found in other closely related strains (Fig. 1), including genes encoding glycosyl hydrolase enzymes with previously reported antibiofilm activity. These findings suggest that purposeful probiotic screening strategies accompanied with comparative genomic analyses can be used to identify unique strains with desired mechanisms of action. This systems approach can further help to elucidate the mechanisms by which a probiotic strain performs a desired action. For instance, a variety of probiotic strains can help resolve *S. aureus* infections through bactericidal mechanisms^8,29^; however, recent evidence has demonstrated certain *Bacillus* strains can also inhibit virulence without killing these pathogenic microorganisms^85,86^. In line with these findings, application of *B. subtilis* 6D1 CFE inhibited *S. aureus* biofilm growth and eradicated mature biofilm in a dose dependent manner (Fig. 3C-D) without inhibiting planktonic growth (Fig. 2A), providing evidence that compounds produced by this strain were capable of quorum sensing interference^87^. Moreover, the ability of *B. subtilis* 6D1 to outcompete *S. aureus* ATCC 29213 in a biofilm but not in a planktonic environment suggest that the antibiofilm activity of this strain is maintained in a coculture environment (Fig. 2C). While we acknowledge these competitive benefits observed in *B. subtilis* 6D1 may be diminished or rendered ineffective in a more complex polymicrobial environment, others report that these beneficial probiotic effects may indeed persist in the human gut and sites of *S. aureus* infection^86,88^. In fact, *B. subtilis* 6D1’s inability to inhibit *S. aureus* planktonic growth was also observed in *B. subtilis* H28, which otherwise successfully alleviated *S. aureus*-induced mastitis infection in a murine model^86^.

Subsequent gene expression assays confirmed application of either *B. subtilis* 6D1 cells or CFE both upregulated genes located in the *Staphylococcus* Agr quorum sensing operon (Fig. 4), further indicating this strain can deploy QSI-mediated antibiofilm mechanisms when in direct competition with *S. aureus*. Specifically, these experiments showed both *B. subtilis* 6D1 culture and its associated CFE increased expression of the same Agr and stress specific genes in *S. aureus* after 24-hour exposure (Fig. 4A). In addition to upregulating *agrA*, expression of both *RNAIII* and *hld* indicate the phosphorylation of AgrA is prompting the transcription of both the P2 and P3 promoters of the Agr regulon (Fig. 4B). Recent work demonstrated that overexpression of *RNAIII* contributes to cell lethality in Agr-positive *S. aureus* strains^89^, suggesting *B. subtilis* 6D1 might induce cell death indirectly if exposure is increased beyond 24 hours and *RNAIII* expression increases. In addition to upregulating *agrA*, *RNAIII*, and *hld*, application of *B. subtilis* 6D1 and its associated CFE also increased the expression of *sigB* and *saeR*. Interestingly, both SigB and the SaeR/S two-component system are essential to persist intracellularly and evade the host innate immune system^90,91^. However, to accomplish this state of persistence, *S. aureus* also needs to silence Agr, and, in doing so, adopt a biofilm lifestyle^75^. The Sae-regulon also includes genes associated with biofilm formation (nucleases) and dispersal (proteases) factors. Therefore, it is possible different compounds produced by *B. subtilis* 6D1 might promote expression of the Sae system independently of the Agr system and negatively impact biofilm formation by stimulating *S. aureus* protease production^70^. In support of this hypothesis, it was recently demonstrated that cyclic dipeptides produced by *Lactobacillus reuteri* RC14 can independently interfere with both the Agr and SaeR/S signaling systems^92^.

Though we were surprised to observe similar *S. aureus* gene expression responses after exposure to *B. subtilis* 6D1 CFE and live cells, these data suggest *B. subtilis* 6D1 and its associated CFE might both inhibit virulence strategies used by pathogenic Staphylococcal species. Furthermore, our data provide additional evidence towards the benefit of using a mixture of probiotic-derived compounds rather than a single compound. The synergistic effects of *B. subtilis* 6D1 CFE when applied alongside low doses of gentamicin and ampicillin (Fig. 3E-F) aligns with previous studies investigating the improved inhibitory effects seen when *B. subtilis* CFE was paired with gentamicin and penicillin to treat an osteomyelitis infection in mice^93^. The overuse of antibiotics is a major concern in both agriculture and hospital settings, and the use of probiotic-derived compounds alongside lower doses of antibiotics may help to reduce the use of antibiotics and decrease the dissemination of antibiotic resistance genes in these environments. Furthermore, application of multiple peptide compounds may also prolong the utility of these therapies by reducing the rate at which resistant mutants appear^94^. In further support of deploying a mixture of probiotic-derived compounds, separation of *B. subtilis* 6D1 CFE into the active fraction 100%P1, comprised of surfactin A-D and an unknown compound, revealed this mixture of compounds harbored greater and broader antibiofilm activity than that of commercial surfactin also obtained from *B. subtilis* (Fig. 6A-E). To our knowledge this is the first study demonstrating *B. subtilis* derived peptides exhibit broad antibiofilm activity against all four *S. aureus* Agr backgrounds. Furthermore, antibiofilm activity against a strain unable to produce AgrD, RN7206, suggest these *B. subtilis* derived peptides are binding AgrC and initiating the phosphorylation of AgrA (Fig. 4B), however more biochemical analyses are required to confirm this hypothesis. Antibiofilm activity of 100%P1 against multiple *S. epidermidis* strains was also somewhat surprising (Fig. 6E)*. S. epidermidis* also harbors an Agr system that succumbs to QSI and cross-talk between similarly structured AgrD configurations^95^, however, the AgrD peptide variants found in *S. epidermidis* are structurally different than those found in *S. aureus*^96^. Despite this, it has been reported that peptides capable of interfering with the *S. aureus* Agr system are also capable of interfering with the *S. epidermidis* Agr systems^56^. This suggests the mixture of peptides produced by *B. subtilis* 6D1 may in fact possess broader QSI activity than tested here. Future studies using purified *B. subtilis* 6D1 surfactins rather than refined fractions will help us to understand the binding affinity and specificity of these compounds to a variety of AgrC receptors found in multiple Staphylococcal Agr signaling systems.

Based on these peptides ability to decrease biofilm formation in multiple Agr backgrounds, we hypothesized that *B. subtilis* 6D1’s ability to reduce biofilm formation by upregulating Agr, may inadvertently increase the expression and production of multiple virulence factors also under control of the Agr system^14^. However, our cell culture data contradicted this hypothesis. The protective and immunomodulatory effects afforded by *B. subtilis* 6D1 CFE in both Vero and HT29 cells indicate the mixture of small peptides produced by this strain can reduce *S. aureus* 29213 virulence through stimulation of intestinal adaptive immunity. One potential explanation for these findings centers around increased transcription of *sigB* (Fig. 4A). Upregulation of SigB has been shown to reduce expression of multiple virulence factors in an effort to subvert the human immune system^91^. However, agr-induced RNAIII levels are elevated in *sigB* mutants^97^, implying that *sigB* expression should be reduced if *agrA* expression increases. This assumption contradicts our findings; however, we hypothesize this difference can be attributed to different CFE compounds independently altering *S. aureus* gene expression associated with biofilm formation and the global stress response. For instance, subtilosin A produced by *Bacillus subtilis* KATMIRA1933 can inhibit biofilm formation in clinical *S. aureus* strains and disrupt quorum sensing in *Listeria monocytogenes* and *Salmonella enterica*^85,98,99^. A biosynthetic gene cluster for subtilosin A was also identified in the *B. subtilis* 6D1 genome with high sequence similarity to the reference cluster (Fig. S3). Therefore, it is possible subtilosin A may be contributing to the QSI or anti virulence activity observed in experiments where crude CFE was employed. This also suggests QSI strategies that induce rather than inhibit the Agr system may also mitigate the virulence potential of *S. aureus*. Furthermore, it is tempting to speculate that QSI strategies employed by probiotic bacteria that facilitate an Agr ‘on’ state might also inadvertently reduce the risk of chronic *S. aureus* infections by separately upregulating genes typically repressed during chronic disease states^16,17^. Interestingly, HT29 monolayers treated with 20%v/v CFE harbored less LDH accumulation than untreated monolayers not challenged with *S. aureus* (data not shown), suggesting this consortium may improve gut epithelial integrity in unchallenged cell types. HT29 cells have been frequently used to study the intestinal immune response to bacterial infection, adhesion, and survival^100^, and although they represent a valuable model due to their similarities with enterocytes of the small intestine, their limitations and the relevance to the *in vivo* situation are still under debate. Specifically, differentiated HT29 cells express brush-border-associated hydrolases typical of the small intestine, however the enzymatic activity is lower than that found *in vivo*. Nevertheless, our results align with previous research demonstrating how probiotic derived small molecules, termed postbiotics, can improve gut physiological processes and adaptive immunity^101,102^.

In addition to exhibiting stronger antibiofilm activity, our cell culture results also demonstrated the 100%P1 fraction was less cytotoxic than commercial surfactin (Fig. 7C) and decreased *S. aureus* induced cytotoxicity in Vero cells when applied at 250 μg/mL (Fig. 7B). Each surfactin isoform (A-D) reduces macrophage cell viability at high concentrations but protects macrophage cells against an LPS challenge at lower concentrations^103^. These data suggest commercial surfactin may have been applied at too high a concentration to observe any protective benefits in Vero cells infected with *S. aureus*. It is also possible that differences between the commercial surfactin product and fraction 100% P1 are due to the combination and varying concentrations of multiple surfactin isoforms in fraction 100%P1 compared to the commercial surfactin that is enriched with surfactin C. Alternatively, this activity may be due to the presence of the uncharacterized compound found in 100%P1, since refined fractions enriched with this compound also inhibited *S. aureus* biofilm growth and disassembled mature biofilm (Fig. S5). In an attempt to purify this compound and elucidate its structure, batch-culturing was used to increase the abundance of this unknown compound, however the resulting concentrations were too low to deduce an accurate molecular formula or structure. We further attempted to enrich this unknown compound by knocking out the *B. subtilis* 6D1 surfactin gene cluster and obtaining new extracts without any surfactins. However, attempts to knockout the surfactin gene cluster using natural transformation, SPP1 phage transduction, and transposon insertion mutagenesis^104^ were all unsuccessful. Further genomic analysis revealed *B. subtilis* 6D1 is missing *comI* - a pBS32 plasmid encoded gene believed to inhibit natural competence in *B. subtilis*^64,105^. Despite identifying an endogenous pBS32-like plasmid in *B. subtilis* 6D1 (Fig. 1B), blast analysis in both the chromosome and plasmid sequence yielded no hits for the *comI* gene, suggesting this strain may have an impaired machinery to uptake foreign DNA.

Collectively, our work demonstrates that *B. subtilis* 6D1 may serve multiple roles in preventing chronic *S. aureus* infection by inhibiting biofilm formation and attachment, increasing sensitivity to antibiotics, and stimulating an adaptive immune response capable of reducing *S. aureus* cytotoxicity and improving intestinal health. A thorough understanding of probiotic strain mechanisms, like those described for *B. subtilis* 6D1, is essential if they are to serve as preventive and/or therapeutic options against biofilm-forming pathogens like *S. aureus*.

## Author Contributions

K.R.H and K.R.L. conceptualized the study. C.R.C. contributed to the study design. K.R.L., D.S.M., W.S., and G.V. conducted the investigation and performed laboratory analysis. K.R.L. performed the data analysis and its visualization and wrote the original draft. All authors helped interpreting the results, edited, and approved the final manuscript.

The authors declare no competing interests.

## Acknowledgements

This work was funded, in part, by the Department of Defense Office of Army Research Grant number W9132T2220001. We would like to thank Nick Konopek, and Grace Schmaling for their laboratory assistance and Dr. Reed Stubbendieck and Dr. Daniel Kearns for their technical expertise and thoughtful discussion. We would also like to thank Dr. Reed Stubbendieck, Dr. Erik Munson, and Dr. Richard Novick for kindly providing clinical and Agr specific staphylococcal strains. We would also like to acknowledge Microbial Discovery Group for their continued support of student research.

