## Supplementary Material for "*Bacillus subtilis* derived lipopeptides disrupt quorum sensing and biofilm assembly in *Staphylococcus aureus*"

### Supplementary Tables

**Table S1. Bacterial Strain List**

| **Strain Name** | **Reference** |
|---|---|
| *S. aureus* ATCC 29213 | American Type Culture Collection |
| *S. aureus* RN6734 (Prototypical agr-I) | Ji et al. 1997 |
| *S. aureus* RN6607 (Prototypical agr-II) | Ji et al. 1997 |
| *S. aureus* RN3984 (Prototypical agr-III) | Ji et al. 1997 |
| *S. aureus* RN4850 (Prototypical agr-IV) | Ji et al. 1997 |
| *S. aureus* RN7206 (agr-null derivative of RN6734) | Ji et al. 1997 |
| *S. aureus* RSM5 | This Paper |
| *S. aureus* RSM97 | This Paper |
| *S. aureus* RSM256 | This Paper |
| *S. aureus* RSM424 | This Paper |
| *S. aureus* RSM486 | This Paper |
| *S. epidermidis* ATCC 12228 | American Type Culture Collection |
| *S. epidermidis* RSM78 | This Paper |
| *S. epidermidis* RSM133 | This Paper |
| *S. epidermidis* RSM242 | This Paper |
| *S. epidermidis* RSM268 | This Paper |
| *S. epidermidis* RSM425 | This Paper |
| *B. subtilis* NCIB 3610 | Bacillus Genetic Stock Center |
| *B. subtilis* 9B5 | This Paper |
| *B. subtilis* 6D1 | This Paper |
| *B. pumilus* 11D12 | This Study |
| *B. pumilus* 10D6 | This Study |
| *B. pumilus* 11G1 | This Study |
| *B. pumilus* 11F9 | This Study |
| *B. pumilus* 11B2 | This Study |
| *Bacillus* 11A10 | This Study |
| *Bacillus* 11A12 | This Study |
| *Bacillus* 3A3 | This Study |
| *Bacillus* 10C11 | This Study |
| *Bacillus* 2G11 | This Study |
| *Bacillus* 11G5 | This Study |
| *Bacillus* 3H6 | This Study |
| *Bacillus* 1E1 | This Study |
| *Bacillus* 5A12 | This Study |

Research supported by the Office of Army Research. The views expressed in this paper are those of the authors and do not reflect the official policy or position of the Department of the Army, Department of Defense, or the U.S. Government.

**Table S2. List of gene expression targets and primer melting temperatures (Tm)**

| Gene | Forward primer (5'-3') | Reverse primer (5'-3') | Tm |
|---|---|---|---|
| *agrA* | TGCGAAGACGATCCAAAAC | TTTAGCTTGCTCAAGCACCTC | 60°C |
| *aur* | GATGGTCGCACATTCACAAG | CGCCTGACTGGTCCTTATATTC | 60°C |
| *lrgB* | TATTGCCCGAGGATTAGCAC | CAAAGACAGGCACAACTGCTAC | 60°C |
| *lukD* | GTACTTAAGGCAGCCGGAAAC | CGCCCCAATAAAACTGTGAG | 60°C |
| *sigB* | TGATCGCGAACGAGAAATC | ATTGCCGTTCTCTGAAGTCG | 60°C |
| *capC* | CATCCAGAGCGGAATAAAGC | CGGAAATACCCGCTAATGAC | 59.5°C |
| *isaA* | TCCGACAAACACTGTTGACC | AATCCCCAAGCACCTAAACC | 59.5°C |
| *lytM* | ACGGTGTCGACTATGCAATG | ATTGCCGCCACCATAGTTAC | 59.5°C |
| *saeR* | CCAAGGGAACTCGTTTTACG | ACGCATAGGGACTTCGTGAC | 59.5°C |
| *fnbB* | GAACATGGTCAAGCACAAGG | ACGCCATAATTACCGTGACC | 59°C |
| *hla* | TCTTGGAACCCGGTATATGG | AGCGAAGTCTGGTGAAAACC | 59°C |
| *hlb* | GTGCCAAAGCCGAATCTAAG | ATCAGCGCGTTTATATTGTCC | 59°C |
| *hld* | AAGGAAGGAGTGATTTCAATGG | TTTGTTCACTGTGTCGATAATCC | 59°C |
| *psmα* | TCAAAAGCTTAATCGAACAATTCAC | AATGGCCCCCTTCAAATAAG | 59°C |
| *sarA* | TTGCTTTGAGTTGTTATCAATGG | CAATACAGCGAATTCTTCAAAGC | 59°C |
| *cidA* | CTTAGCCGGCAGTATTGTTG | GTTTGCACCGTCTTCTACCC | 58.5°C |
| *clfB* | TTATGGTGGTGGAAGTGCTG | TGGACTTGGTTCTGGATCTG | 58.5°C |
| *RNAIII* | AAGCCATCCCAACTTAATAACC | GCACTGAGTCCAAGGAAACTAAC | 58.5°C |
| *rpoB* | ACAACCACTTGGCGGTAAAG | ATGCTTCAAGTGCCCATACC | 60°C |

Research supported by the Office of Army Research. The views expressed in this paper are those of the authors and do not reflect the official policy or position of the Department of the Army, Department of Defense, or the U.S. Government.

#### Table S3. List of Unique Genes Identified in *B. subtilis* 6D1 Genome

| Description | Seed ortholog | e-value | Max annotation level | COG category |
|---|---|---|---|---|
| Prophage endopeptidase tail | 279010.BL03515 | 1.9E-275 | Bacilli | Cell motility; Intracellular trafficking & secretion |
| Catalyzes the first step in the D-alanylation of lipoteichoic acid | 224308.BSU18320 | 0 | Bacilli | Secondary structure |
| Peptidase activity | 10717.Q9ZXF7_BPPH1 | 1.3E-102 | Siphoviridae | Function unknown |
| Phage terminase small subunit | 1274524.BSONL12_06108 | 2.1E-101 | Bacilli | Replication and repair |
| Protein of unknown function (DUF669) | 279010.BL03492 | 1.67E-93 | Bacilli | Function unknown |
| Phosphoribosyl-ATP pyrophosphohydrolase | 1408303.JNJJ01000009_gene2204 | 6.13E-53 | Bacilli | Function unknown |
| Sequence-specific DNA binding | 10717.Q786F1_BPPH1 | 2.01E-82 | Siphoviridae | Function unknown |
| Phage head-tail joining protein | 1274524.BSONL12_06148 | 6.1E-51 | Bacilli | Function unknown |
| BhlA holin family | 224308.BSU21420 | 4.44E-38 | Bacilli | Function unknown |
| Phage tail protein | 279010.BL03514 | 1.5E-131 | Bacilli | Function unknown |
| DNA gyrase B | 881953.E0YIY1_9CAUD | 4.7E-228 | Caudovirales | Function unknown |
| Nuclease activity | 10717.Q9ZXC2_BPPH1 | 3.1E-121 | Siphoviridae | Function unknown |
| Phage portal protein | 1434319.W5RVB1_9CAUD | 1.5E-130 | Siphoviridae | Function unknown |
| Phage portal protein | 66692.ABC1331 | 1.1E-219 | Bacilli | Function unknown |
| DNA gyrase/topoisomerase IV, subunit A | 881953.E0YIY2_9CAUD | 9.4E-181 | Caudovirales | Function unknown |
| Major facilitator superfamily | 1348908.KI518625_gene1647 | 4.37E-26 | Bacilli | Amino Acid/carbohydrate/inorganic ion metabolism & transport |
| Recombinase | 10717.Q9T200_BPPH1 | 4E-288 | Siphoviridae | Function unknown |
| HNH endonuclease | 1274524.BSONL12_06103 | 2.8E-65 | Bacilli | Defense mechanism |
| Nucleoside 2-deoxyribosyltransferase | 1384057.CD33_16495 | 5.17E-62 | Bacilli | Nucleotide metabolism and transport |
| PemK-like, MazF-like toxin of type II toxin-antitoxin system | 1297581.H919_00930 | 4.08E-18 | Bacilli | Signal transduction |
| SPP1 phage holin | 240302.BN982_00173 | 7.04E-14 | Bacilli | Function unknown |
| capsid protein | 66692.ABC1333 | 1.9E-204 | Bacilli | Function unknown |
| Terminase | 1274524.BSONL12_06113 | 0 | Bacilli | Function unknown |
| Guanylate kinase homologues. | 596330.HMPREF0628_1632 | 2.92E-09 | Clostridia | Nucleotide metabolism and transport |
| DNA packaging | 224308.BSU26200 | 3.62E-59 | Bacilli | Replication and repair |
| Phage tail tube protein | 1196324.A374_08734 | 1.08E-35 | Bacilli | Function unknown |

Research supported by the Office of Army Research. The views expressed in this paper are those of the authors and do not reflect the official policy or position of the Department of the Army, Department of Defense, or the U.S. Government.

| | | | | |
|---|---|---|---|---|
| Hydrolase activity | 10717.Q9ZXC4_BPPH1 | 0 | Siphoviridae | Function unknown |
| DNA protection | 10717.Q9ZXC8_BPPH1 | 3.1E-119 | Siphoviridae | Function unknown |
| Caudovirus prohead serine protease | 240302.BN982_03592 | 4.5E-65 | Bacilli | Function unknown |
| N-acetylmuramoyl-L-alanine amidase activity | 1406785.U5PU90_9CAUD | 1.38E-85 | Caudovirales | Function unknown |
| DNA packaging | 1033739.CAEU01000069_gene8 | 4.26E-29 | Bacteria | Function unknown |
| Phage tail tape measure protein | 1274524.BSONL12_06173 | 0 | Bacilli | Cell cycle control and mitosis |
| AAA domain | 1178540.BA70_16395 | 7.5E-211 | Bacilli | Cell cycle control and mitosis |
| 3D domain | 66692.ABC0489 | 1.51E-29 | Bacilli | Cell membrane biogenesis |
| DNA primase activity | 1434319.W5RVC4_9CAUD | 1.8E-113 | Siphoviridae | Function unknown |
| Phage tail tape measure protein | 1340434.AXVA01000005_gene4862 | 2.1E-251 | Bacilli | Cell cycle control and mitosis |
| Phage integrase family | 1434319.W5RV39_9CAUD | 3.7E-122 | Siphoviridae | Function unknown |
| Ribonucleotide reductase, small chain | 1434319.W5RVC6_9CAUD | 7.5E-130 | Siphoviridae | Function unknown |
| Phage tail protein | 411465.PEPMIC_00022 | 2.3E-239 | Firmicutes | Function unknown |
| HNH endonuclease | 1500386.A0A060AKD6_9CAUD | 3.64E-41 | Siphoviridae | Function unknown |
| Pfam:Peptidase_M78 | 10717.Q9T201_BPPH1 | 4.72E-92 | Siphoviridae | Function unknown |
| Helix-turn-helix | 86416.Clopa_0552 | 1.62E-12 | Clostridia | Transcription |
| Aspartate phosphatase | 720555.BATR1942_06825 | 5.4E-135 | Bacilli | Function unknown |
| Phage terminase, small subunit | 1500386.A0A060AKR4_9CAUD | 4.92E-39 | Siphoviridae | Function unknown |
| Immunity protein 70 | 1274524.BSONL12_12931 | 3.62E-86 | Bacilli | Function unknown |
| Immunity protein 70 | 1274524.BSONL12_06218 | 2.6E-82 | Bacilli | Function unknown |
| Phage terminase | 1033739.CAEU01000068_gene122 | 3.4E-261 | Bacilli | Function unknown |
| DNA helicase activity | 1500386.A0A060AB93_9CAUD | 1.6E-132 | Siphoviridae | Function unknown |
| Glycosyl hydrolases family 25 | 666686.B1NLA3E_01320 | 1E-145 | Bacilli | Cell membrane biogenesis |
| Transcriptional regulator | 265729.GS18_0220175 | 4.38E-13 | Bacilli | Transcription |
| Bacteriophage HK97-gp10, putative tail-component | 1274524.BSONL12_06153 | 2.81E-65 | Bacilli | Function unknown |
| YopX protein | 1536769.P40081_15180 | 4.03E-11 | Bacilli | Function unknown |
| PfkB family carbohydrate kinase | 272556.CF65_00429 | 3.71E-09 | Gamma-proteobacteria | Carbohydrate metabolism and transport |
| Replication initiator protein A (RepA) N-terminus | 1329250.WOSG25_090160 | 1.57E-15 | Bacilli | Function unknown |

Research supported by the Office of Army Research. The views expressed in this paper are those of the authors and do not reflect the official policy or position of the Department of the Army, Department of Defense, or the U.S. Government.

| | | | | |
|---|---|---|---|---|
| Endonuclease that specifically degrades the RNA of RNA- DNA hybrids | 1128398.Curi_c12400 | 4.61E-60 | Clostridia | Replication and repair |
| Uncharacterized protein YqaH | 1178540.BA70_16380 | 1.78E-28 | Bacilli | Function unknown |
| Phage gp6-like head-tail connector protein | 1274524.BSONL12_06143 | 8.77E-54 | Bacilli | Function unknown |
| Phage tail tube protein | 1274524.BSONL12_06163 | 3.2E-101 | Bacilli | Function unknown |
| Domain of unknown function (DUF1738) | 1415775.U729_3188 | 1.4E-71 | Clostridia | Replication and repair |
| Phage capsid family | 1034347.CAHJ01000091_gene1634 | 1.6E-108 | Bacilli | Function unknown |
| Integrase family | 1321778.HMPREF1982_00376 | 1.21E-92 | Clostridia | Replication and repair |
| Ribonucleoside-diphosphate reductase activity, thioredoxin disulfide as acceptor | 1434319.W5RV23_9CAUD | 0 | Siphoviridae | Function unknown |
| Bacterial DNA polymerase III alpha subunit | 1434319.W5RV95_9CAUD | 0 | Siphoviridae | Function unknown |
| Flavodoxin | 1444310.JANV01000144_gene363 | 6.76E-12 | Bacilli | Energy production and conversion |

**Table S4. List of Single Copy Orthologs Used for *B. subtilis* 6D1 Phylogenetic Analysis**

| PGFam | Alignment Score | Alignment Length | Mean Square Frequency | Proposed Number of Gaps | Product |
|---|---|---|---|---|---|
| PGF_02704551 | 34.62 | 1199 | 1 | 0 | DNA-directed RNA polymerase beta' subunit (EC 2.7.7.6) |
| PGF_00045963 | 33.84 | 1148 | 0.999 | 0 | Pyruvate carboxylase (EC 6.4.1.1) |
| PGF_07830674 | 29.51 | 878 | 0.996 | 0 | Alanyl-tRNA synthetase (EC 6.1.1.7) |
| PGF_07058357 | 27.93 | 786 | 0.996 | 0 | Single-stranded-DNA-specific exonuclease RecJ |
| PGF_04438983 | 27.81 | 774 | 1 | 0 | ATP-dependent protease La (EC 3.4.21.53) Type I |
| PGF_12779560 | 27.46 | 775 | 0.986 | 0 | Phage infection protein |
| PGF_01444054 | 27.28 | 749 | 0.997 | 0 | ATP-dependent RNA helicase |
| PGF_01671660 | 26.69 | 801 | 0.943 | 0.046 | Transcriptional regulator, AraC family |
| PGF_04883561 | 26.28 | 710 | 0.986 | 0 | Assimilatory nitrate reductase large subunit (EC 1.7.99.4) |
| PGF_00840774 | 25.82 | 673 | 0.995 | 0 | Heterodimeric efflux ABC transporter, multidrug resistance => BmrD subunit of BmrCD |
| PGF_06875694 | 25.76 | 668 | 0.997 | 0 | Penicillin-binding protein 3 |
| PGF_04333086 | 25.73 | 668 | 0.995 | 0 | DNA ligase (NAD(+)) (EC 6.5.1.2) |
| PGF_06459503 | 25.41 | 648 | 0.998 | 0 | Serine/threonine protein kinase PrkC, regulator of stationary phase |
| PGF_00005140 | 25.32 | 646 | 0.996 | 0 | ABC transporter-like sensor and permease protein YvcS |
| PGF_00960048 | 25.04 | 639 | 0.991 | 0.006 | PTS system, fructose-specific IIA component (EC 2.7.1.202) / PTS system, fructose-specific IIB component (EC |

Research supported by the Office of Army Research. The views expressed in this paper are those of the authors and do not reflect the official policy or position of the Department of the Army, Department of Defense, or the U.S. Government.

| | | | | | |
|---|---|---|---|---|---|
| | | | | | 2.7.1.202) / PTS system, fructose-specific IIC component |
| PGF_06841555 | 24.93 | 643 | 0.983 | 0.001 | hypothetical protein |
| PGF_07573550 | 24.78 | 622 | 0.993 | 0 | ABC transporter-like sensor and permease protein YxdM |
| PGF_02788357 | 24.66 | 611 | 0.998 | 0.001 | Two-component sensor kinase SA14-24 |
| PGF_00012969 | 24.21 | 590 | 0.997 | 0 | Phosphomethylpyrimidine synthase ThiC (EC 4.1.99.17) |
| PGF_00045994 | 24.16 | 585 | 0.999 | 0 | Pyruvate kinase (EC 2.7.1.40) / Phosphohistidine swiveling domain |
| PGF_00066435 | 24.12 | 643 | 0.951 | 0.045 | Asparagine synthetase [glutamine-hydrolyzing] (EC 6.3.5.4) YisO |
| PGF_10523783 | 23.89 | 572 | 0.999 | 0 | Acetyl-CoA synthetase (EC 6.2.1.1) |
| PGF_00025227 | 23.87 | 571 | 0.999 | 0 | Acetolactate synthase, catabolic (EC 2.2.1.6) |
| PGF_00055052 | 23.85 | 571 | 0.998 | 0 | Sulfite reductase [NADPH] hemoprotein beta-component (EC 1.8.1.2) |
| PGF_08632970 | 23.8 | 573 | 0.994 | 0 | diguanylate cyclase/phosphodiesterase (GGDEF & EAL domains) with PAS/PAC sensor(s) |
| PGF_00421624 | 23.7 | 577 | 0.987 | 0.012 | DNA polymerase X family |
| PGF_00051108 | 23.59 | 562 | 0.995 | 0.001 | Septation ring formation regulator EzrA |
| PGF_00990926 | 23.43 | 562 | 0.988 | 0.001 | Trehalose-6-phosphate hydrolase (EC 3.2.1.93) |
| PGF_00419557 | 23.19 | 541 | 0.997 | 0 | Copper resistance protein CopC / Copper resistance protein CopD |
| PGF_00423553 | 23.08 | 549 | 0.985 | 0.006 | Dipeptide ABC transporter, substrate-binding protein DppA (TC 3.A.1.5.2) |
| PGF_03294256 | 22.68 | 536 | 0.979 | 0.017 | PTS system, maltose-specific IIC component / PTS system, maltose-specific IIB component (EC 2.7.1.208) |
| PGF_04294677 | 22.56 | 518 | 0.991 | 0.002 | 2-isopropylmalate synthase (EC 2.3.3.13) |
| PGF_07491427 | 22.52 | 509 | 0.998 | 0 | Inner membrane protein YqiK |
| PGF_00056814 | 22.49 | 515 | 0.991 | 0 | Anthranilate synthase, aminase component (EC 4.1.3.27) |
| PGF_00013509 | 22.49 | 512 | 0.994 | 0 | IMP cyclohydrolase (EC 3.5.4.10) / Phosphoribosylaminoimidazolecarboxamide formyltransferase (EC 2.1.2.3) |
| PGF_01675349 | 22.48 | 510 | 0.996 | 0 | Purine nucleoside ABC transporter, ATP-binding protein |
| PGF_00013418 | 22.32 | 501 | 0.997 | 0 | Oxygen-independent coproporphyrinogen-III oxidase-like protein HemZ |
| PGF_00420806 | 22.31 | 503 | 0.995 | 0 | D-alanine--poly(phosphoribitol) ligase subunit 1 (EC 6.1.1.13) |
| PGF_00007119 | 22.25 | 513 | 0.982 | 0.001 | Galactose-1-phosphate uridylyltransferase (EC 2.7.7.10) |
| PGF_00054087 | 22.18 | 492 | 1 | 0 | Stage IV sporulation protein A |
| PGF_05424184 | 22.11 | 496 | 0.993 | 0 | Na(+) H(+) antiporter subunit D |
| PGF_00402817 | 22.09 | 490 | 0.998 | 0 | Betaine aldehyde dehydrogenase (EC 1.2.1.8) |
| PGF_00962420 | 22.08 | 490 | 0.998 | 0 | Arginine decarboxylase (EC 4.1.1.19) |

Research supported by the Office of Army Research. The views expressed in this paper are those of the authors and do not reflect the official policy or position of the Department of the Army, Department of Defense, or the U.S. Government.

| | | | | | |
|---|---|---|---|---|---|
| PGF_00008337 | 21.94 | 483 | 0.998 | 0 | Glutamyl-tRNA synthetase (EC 6.1.1.17) @ Glutamyl-tRNA(Gln) synthetase (EC 6.1.1.24) |
| PGF_10345122 | 21.81 | 479 | 0.997 | 0 | Lactate utilization protein LutB |
| PGF_00066854 | 21.76 | 475 | 0.998 | 0 | Aspartate ammonia-lyase (EC 4.3.1.1) |
| PGF_00956915 | 21.74 | 486 | 0.986 | 0 | O-succinylbenzoic acid--CoA ligase (EC 6.2.1.26) |
| PGF_00008876 | 21.7 | 484 | 0.987 | 0 | Glycogen synthase, ADP-glucose transglucosylase (EC 2.4.1.21) |
| PGF_00012901 | 21.69 | 473 | 0.997 | 0 | Hydroxyaromatic non-oxidative decarboxylase protein C (EC 4.1.1.-) |
| PGF_03068639 | 21.54 | 471 | 0.993 | 0.002 | Amino-acid permease RocC |
| PGF_03446029 | 21.54 | 475 | 0.988 | 0.01 | PTS system, trehalose-specific IIB component (EC 2.7.1.201) / PTS system, trehalose-specific IIC component |
| PGF_06132116 | 21.49 | 463 | 0.999 | 0 | L-cystine uptake protein TcyP, sodium:anion symporter family |
| PGF_00035888 | 21.45 | 470 | 0.989 | 0 | Predicted glycolate dehydrogenase, 2-subunit type (EC 1.1.99.14), iron-sulfur subunit GlcD |
| PGF_00064101 | 21.41 | 464 | 0.994 | 0.001 | UDP-glucose dehydrogenase in teichuronic acid synthesis TuaD (EC 1.1.1.22) |
| PGF_02620298 | 21.37 | 461 | 0.995 | 0 | Argininosuccinate lyase (EC 4.3.2.1) |
| PGF_10483952 | 21.31 | 459 | 0.995 | 0 | PTS system, IIB component / PTS system, IIC component |
| PGF_00064600 | 21.24 | 475 | 0.975 | 0.02 | Uncharacterized RNA methyltransferase YfjO |
| PGF_02648683 | 21.24 | 456 | 0.995 | 0.002 | PTS system, N-acetylmuramic acid-specific IIB component (EC 2.7.1.192) / PTS system, N-acetylmuramic acid-specific IIC component |
| PGF_03405430 | 21.24 | 455 | 0.996 | 0 | D-glucarate transporter |
| PGF_03318742 | 21.23 | 452 | 0.999 | 0 | PTS system, N-acetylglucosamine-specific IIC component / PTS system, N-acetylglucosamine-specific IIB component (EC 2.7.1.193) |
| PGF_04560429 | 21.23 | 451 | 1 | 0 | Mg/Co/Ni transporter MgtE, CBS domain-containing |
| PGF_00053892 | 21.18 | 450 | 0.998 | 0 | Spore germination protein YpeB |
| PGF_00066263 | 21.13 | 479 | 0.966 | 0.019 | Uronate isomerase (EC 5.3.1.12) |
| PGF_00880747 | 21.12 | 453 | 0.992 | 0 | Na+/H+ antiporter NhaC |
| PGF_00008774 | 21.11 | 448 | 0.997 | 0 | Glycine dehydrogenase [decarboxylating] (glycine cleavage system P1 protein) (EC 1.4.4.2) |
| PGF_00184655 | 21.09 | 448 | 0.997 | 0 | hypothetical protein |
| PGF_00053945 | 21.05 | 445 | 0.998 | 0 | Sporulation protein YkvU |
| PGF_00769755 | 21.05 | 447 | 0.995 | 0 | 16S rRNA (cytosine(967)-C(5))-methyltransferase (EC 2.1.1.176) |
| PGF_03146251 | 20.91 | 450 | 0.985 | 0.002 | Uric acid permease PucJ |
| PGF_00028525 | 20.89 | 441 | 0.995 | 0.004 | Oxaloacetate decarboxylase involved in citrate fermentation (EC 4.1.1.3) |

Research supported by the Office of Army Research. The views expressed in this paper are those of the authors and do not reflect the official policy or position of the Department of the Army, Department of Defense, or the U.S. Government.

| | | | | | |
|---|---|---|---|---|---|
| PGF_00007024 | 20.84 | 436 | 0.998 | 0 | GTP-binding protein EngA |
| PGF_00058560 | 20.84 | 521 | 0.913 | 0.08 | Transcriptional regulator GabR of GABA utilization (GntR family with aminotransferase-like domain) |
| PGF_00421032 | 20.81 | 448 | 0.983 | 0 | D-serine ammonia-lyase (EC 4.3.1.18) |
| PGF_00066502 | 20.76 | 431 | 1 | 0 | Asparaginyl-tRNA synthetase (EC 6.1.1.22) |
| PGF_06935032 | 20.73 | 430 | 1 | 0 | Adenylosuccinate synthetase (EC 6.3.4.4) |
| PGF_00064241 | 20.72 | 441 | 0.987 | 0 | UPF0214 protein YfeW |
| PGF_01054379 | 20.71 | 431 | 0.998 | 0 | Adenylosuccinate lyase (EC 4.3.2.2) @ SAICAR lyase (EC 4.3.2.2) |
| PGF_03064306 | 20.68 | 428 | 0.999 | 0 | Sporulation kinase C (EC 2.7.13.3) |
| PGF_00066695 | 20.67 | 443 | 0.982 | 0 | Vitamin B12 ABC transporter, ATP-binding protein BtuD / Adenosylcobinamide amidohydrolase (EC 3.5.1.90) |
| PGF_09188652 | 20.64 | 430 | 0.995 | 0 | Molybdopterin molybdenumtransferase (EC 2.10.1.1) |
| PGF_02939833 | 20.63 | 434 | 0.99 | 0.001 | UPF0053 membrane protein YrkA |
| PGF_00001028 | 20.61 | 428 | 0.996 | 0 | Uncharacterized zinc protease YmfH |
| PGF_06833830 | 20.6 | 436 | 0.987 | 0 | hypothetical protein |
| PGF_02452671 | 20.55 | 424 | 0.998 | 0 | Cell division trigger factor (EC 5.2.1.8) |
| PGF_09398028 | 20.54 | 422 | 1 | 0 | Uncharacterized protease YrrO |
| PGF_00015701 | 20.53 | 423 | 0.998 | 0 | Isocitrate dehydrogenase [NADP] (EC 1.1.1.42) |
| PGF_03004613 | 20.51 | 424 | 0.996 | 0 | Histidyl-tRNA synthetase (EC 6.1.1.21) |
| PGF_03515627 | 20.49 | 426 | 0.993 | 0 | hypothetical protein |
| PGF_03685660 | 20.45 | 428 | 0.988 | 0.002 | Uncharacterized MFS-type transporter YxiO |
| PGF_00049731 | 20.45 | 421 | 0.996 | 0 | Aluminum resistance protein |
| PGF_00066935 | 20.44 | 466 | 0.947 | 0.045 | Uracil permease @ Uracil:proton symporter UraA |
| PGF_01124177 | 20.33 | 417 | 0.996 | 0 | Dihydrolipoamide succinyltransferase component (E2) of 2-oxoglutarate dehydrogenase complex (EC 2.3.1.61) |
| PGF_05387084 | 20.25 | 415 | 0.994 | 0 | Gamma-glutamyl phosphate reductase (EC 1.2.1.41) |
| PGF_01443078 | 20.18 | 409 | 0.998 | 0 | Predicted signal transduction protein |
| PGF_00736716 | 20.15 | 412 | 0.993 | 0 | hypothetical protein |
| PGF_00021976 | 20.06 | 404 | 0.998 | 0 | Uncharacterized metal ion transporter YcsG, Mn(2+)/Fe(2+) NRAMP family |
| PGF_03202156 | 20 | 400 | 1 | 0 | S-adenosylmethionine synthetase (EC 2.5.1.6) |
| PGF_00092388 | 19.94 | 410 | 0.985 | 0.006 | hypothetical protein |
| PGF_00034602 | 19.82 | 393 | 1 | 0 | Poly-gamma-glutamate synthase subunit PgsB/CapB (EC 6.3.2.-) |
| PGF_02944756 | 19.74 | 391 | 0.998 | 0 | 3-ketoacyl-CoA thiolase [fadN-fadA-fadE operon] (EC 2.3.1.16) |
| PGF_00023189 | 19.73 | 397 | 0.99 | 0 | Uncharacterized MFS-type transporter YttB |
| PGF_10555225 | 19.67 | 412 | 0.969 | 0.022 | Ornithine aminotransferase (EC 2.6.1.13) |

Research supported by the Office of Army Research. The views expressed in this paper are those of the authors and do not reflect the official policy or position of the Department of the Army, Department of Defense, or the U.S. Government.

| | | | | | |
|---|---|---|---|---|---|
| PGF_00403927 | 19.67 | 393 | 0.992 | 0 | Biosynthetic Aromatic amino acid aminotransferase alpha (EC 2.6.1.57) @ Aspartate aminotransferase (EC 2.6.1.1) |
| PGF_09155108 | 19.63 | 391 | 0.993 | 0 | Exonuclease SbcD |
| PGF_00054051 | 19.6 | 411 | 0.967 | 0.029 | Stage III sporulation protein AE |
| PGF_00417840 | 19.52 | 396 | 0.981 | 0.016 | Chorismate synthase (EC 4.2.3.5) |
| PGF_00056157 | 19.51 | 389 | 0.989 | 0 | Teichuronic acid biosynthesis glycosyl transferase TuaC |
| PGF_10149521 | 19.47 | 380 | 0.999 | 0 | Glucose-1-phosphate adenylyltransferase (EC 2.7.7.27) |
| PGF_00027514 | 19.46 | 385 | 0.992 | 0 | N-acetylornithine aminotransferase (EC 2.6.1.11) |
| PGF_09438139 | 19.39 | 378 | 0.998 | 0 | Flagellar motor switch protein FliN |
| PGF_00032869 | 19.38 | 380 | 0.994 | 0 | Isovaleryl-CoA dehydrogenase (EC 1.3.8.4) |
| PGF_04991657 | 19.38 | 379 | 0.995 | 0 | Oxygen-independent coproporphyrinogen-III oxidase-like protein YggW |
| PGF_03048863 | 19.37 | 377 | 0.997 | 0 | Response regulator aspartate phosphatase B |
| PGF_03028706 | 19.36 | 377 | 0.997 | 0 | Spore coat protein CotSA |
| PGF_00420020 | 19.35 | 380 | 0.992 | 0.003 | Cystathionine gamma-lyase (EC 4.4.1.1) @ Homocysteine desulfhydrase (EC 4.4.1.2) |
| PGF_00008313 | 19.34 | 415 | 0.949 | 0.042 | Glutamine-dependent 2-keto-4-methylthiobutyrate transaminase |
| PGF_02653253 | 19.28 | 378 | 0.992 | 0.003 | ABC transporter, RND-adapter-like protein YknX |
| PGF_00024274 | 19.27 | 380 | 0.989 | 0 | N5-carboxyaminoimidazole ribonucleotide synthase (EC 6.3.4.18) |
| PGF_07072582 | 19.26 | 371 | 1 | 0 | RNA polymerase sigma factor RpoD |
| PGF_07063065 | 19.26 | 371 | 1 | 0 | Transcription termination protein NusA |
| PGF_00502238 | 19.26 | 371 | 1 | 0 | Pyruvate dehydrogenase E1 component alpha subunit (EC 1.2.4.1) |
| PGF_00024656 | 19.22 | 382 | 0.983 | 0.005 | Uncharacterized NADH-dependent flavin oxidoreductase YqiG |
| PGF_00404036 | 19.21 | 410 | 0.949 | 0.035 | Biotin biosynthesis cytochrome P450 (EC 1.14.15.12) |
| PGF_04871820 | 19.21 | 371 | 0.997 | 0 | Prephenate dehydrogenase (EC 1.3.1.12) |
| PGF_00722427 | 19.18 | 373 | 0.993 | 0 | hypothetical protein |
| PGF_00007012 | 19.11 | 366 | 0.999 | 0 | GTP-binding and nucleic acid-binding protein YchF |
| PGF_00056316 | 19.1 | 368 | 0.996 | 0 | Tetraprenyl-beta-curcumene synthase (EC 4.2.3.130) |
| PGF_00016850 | 19.06 | 364 | 0.999 | 0 | Branched-chain amino acid dehydrogenase [deaminating] (EC 1.4.1.9)(EC 1.4.1.23) |
| PGF_04146410 | 19.05 | 363 | 1 | 0 | Protein-arginine kinase McsB (EC 2.7.14.1) |
| PGF_00408522 | 19.05 | 371 | 0.989 | 0 | Putative isomerase YitF |
| PGF_00007560 | 19.04 | 366 | 0.995 | 0 | Germination (Cortex hydrolysis) and sporulation protein GerM |
| PGF_03961783 | 19.03 | 365 | 0.996 | 0 | Uncharacterized protein YvlB |
| PGF_00754446 | 18.99 | 410 | 0.938 | 0.048 | Uncharacterized MFS-type transporter YceJ |

Research supported by the Office of Army Research. The views expressed in this paper are those of the authors and do not reflect the official policy or position of the Department of the Army, Department of Defense, or the U.S. Government.

| | | | | | |
|---|---|---|---|---|---|
| PGF_09945671 | 18.98 | 433 | 0.912 | 0.085 | Acetate kinase (EC 2.7.2.1) |
| PGF_00037527 | 18.95 | 364 | 0.993 | 0.001 | Proline dipeptidase (EC 3.4.13.9) |
| PGF_00069766 | 18.95 | 365 | 0.992 | 0 | Type III polyketide sythase producing alkylpyrones (*B. subtilis* BpsA) |
| PGF_01756633 | 18.94 | 399 | 0.948 | 0.03 | Ferrichrome transport system permease protein FhuB |
| PGF_00828128 | 18.93 | 361 | 0.996 | 0 | Putative aminopeptidase YsdC |
| PGF_04858171 | 18.93 | 363 | 0.993 | 0.001 | Branched-chain acyl kinase |
| PGF_00014295 | 18.91 | 362 | 0.994 | 0 | Inner spore coat protein CotH |
| PGF_00030640 | 18.87 | 356 | 1 | 0 | Peptide chain release factor 1 |
| PGF_08308923 | 18.86 | 357 | 0.998 | 0 | Sigma-M negative effector |
| PGF_02516666 | 18.83 | 356 | 0.998 | 0 | Cytochrome c oxidase polypeptide II (EC 1.9.3.1) |
| PGF_00054982 | 18.71 | 354 | 0.994 | 0 | Sulfate permease, Pit-type |
| PGF_00053851 | 18.65 | 351 | 0.996 | 0 | Spore coat protein CotS |
| PGF_00028828 | 18.62 | 356 | 0.987 | 0 | Uncharacterized oxidoreductase YxnA |
| PGF_09679949 | 18.57 | 358 | 0.981 | 0.005 | DNA alkylation repair enzyme |
| PGF_07915158 | 18.52 | 346 | 0.996 | 0 | N(6)-L-threonylcarbamoyladenine synthase (EC 2.3.1.234) |
| PGF_00811541 | 18.49 | 347 | 0.993 | 0 | Uncharacterized membrane protein YkvI |
| PGF_00758189 | 18.43 | 344 | 0.994 | 0 | hypothetical protein |
| PGF_07889681 | 18.39 | 345 | 0.99 | 0 | N-acetyl-gamma-glutamyl-phosphate reductase (EC 1.2.1.38) |
| PGF_00247656 | 18.38 | 340 | 0.997 | 0 | Putative membrane-bound acyltransferase YkrP |
| PGF_00417381 | 18.37 | 340 | 0.997 | 0 | Central glycolytic genes regulator |
| PGF_00648450 | 18.36 | 337 | 1 | 0 | Rod shape-determining protein MreB |
| PGF_00008864 | 18.35 | 345 | 0.988 | 0.006 | Glycogen biosynthesis protein GlgD, glucose-1-phosphate adenylyltransferase family |
| PGF_00927902 | 18.33 | 348 | 0.983 | 0.002 | hypothetical protein |
| PGF_00024478 | 18.3 | 335 | 1 | 0 | NAD-dependent glyceraldehyde-3-phosphate dehydrogenase (EC 1.2.1.12) |
| PGF_00025057 | 18.27 | 355 | 0.97 | 0.018 | NTD biosynthesis operon putative oxidoreductase NtdC (EC 1.-.-.-) |
| PGF_00037171 | 18.24 | 333 | 1 | 0 | Probable low-affinity inorganic phosphate transporter |
| PGF_00689473 | 18.2 | 361 | 0.958 | 0.029 | Allergen V5/Tpx-1 related |
| PGF_02944779 | 18.19 | 333 | 0.997 | 0 | Acetoin dehydrogenase E1 component alpha-subunit (EC 2.3.1.190) |
| PGF_03198682 | 18.16 | 335 | 0.992 | 0 | Protease IV |
| PGF_10315399 | 18.15 | 339 | 0.986 | 0 | Oxidoreductase, zinc-binding dehydrogenase family (EC 1.1.1.-) |
| PGF_00907539 | 18.13 | 330 | 0.998 | 0 | Branched-chain alpha-keto acid dehydrogenase, E1 component, alpha subunit (EC 1.2.4.4) |
| PGF_03677192 | 18.13 | 331 | 0.997 | 0 | Lipoate-protein ligase A |

Research supported by the Office of Army Research. The views expressed in this paper are those of the authors and do not reflect the official policy or position of the Department of the Army, Department of Defense, or the U.S. Government.

| | | | | | |
|---|---|---|---|---|---|
| PGF_00008611 | 18.12 | 345 | 0.975 | 0.001 | Glycerol-3-phosphate dehydrogenase [NAD(P)+] (EC 1.1.1.94) |
| PGF_00412879 | 18.09 | 329 | 0.998 | 0 | Putative glycosyltransferase CsbB |
| PGF_00412178 | 18.08 | 327 | 1 | 0 | Branched-chain alpha-keto acid dehydrogenase, E1 component, beta subunit (EC 1.2.4.4) |
| PGF_02881645 | 18.08 | 332 | 0.992 | 0.006 | Transcriptional regulator GanR, LacI family |
| PGF_00070244 | 18.04 | 331 | 0.991 | 0 | BH0638 unknown conserved protein |
| PGF_01975849 | 18.02 | 337 | 0.982 | 0 | (R)-2-hydroxyacid dehydrogenase, similar to L-sulfolactate dehydrogenase (EC 1.1.1.272) |
| PGF_05399159 | 18.01 | 325 | 0.999 | 0 | Acetyl-coenzyme A carboxyl transferase alpha chain (EC 6.4.1.2) |
| PGF_00417233 | 17.93 | 332 | 0.984 | 0.007 | L-Ala--D-Glu endopeptidase |
| PGF_06874967 | 17.89 | 322 | 0.997 | 0 | [4Fe-4S]-AdoMet protein YtqA |
| PGF_06547054 | 17.86 | 322 | 0.996 | 0 | Ribose ABC transporter, permease protein RbsC (TC 3.A.1.2.1) |
| PGF_09753067 | 17.86 | 322 | 0.995 | 0 | Uncharacterized membrane protein YqjA |
| PGF_02614796 | 17.82 | 324 | 0.99 | 0.003 | Uncharacterized membrane protein YkoY |
| PGF_00178344 | 17.79 | 320 | 0.995 | 0.001 | Putative sporulation hydrolase CotR |
| PGF_00869466 | 17.76 | 317 | 0.998 | 0 | Transcriptional activator protein med |
| PGF_00033289 | 17.72 | 340 | 0.961 | 0.029 | Phosphate:acyl-ACP acyltransferase PlsX (EC 2.3.1.n2) |
| PGF_00416436 | 17.7 | 314 | 0.999 | 0.001 | 3'->5' exoribonuclease Bsu YhaM |
| PGF_12795189 | 17.7 | 324 | 0.983 | 0 | Transcriptional regulator, LysR family |
| PGF_02191019 | 17.68 | 325 | 0.981 | 0 | Thiamine-monophosphate kinase (EC 2.7.4.16) |
| PGF_00069761 | 17.68 | 318 | 0.991 | 0.001 | Cephalosporin-C deacetylase (EC 3.1.1.41) |
| PGF_00064754 | 17.67 | 319 | 0.989 | 0.009 | Uncharacterized iron compound ABC uptake transporter, permease protein |
| PGF_10097367 | 17.65 | 314 | 0.996 | 0 | Porphobilinogen deaminase (EC 2.5.1.61) |
| PGF_00064130 | 17.64 | 316 | 0.992 | 0 | UDP-glucuronate 5'-epimerase (EC 5.1.3.12) |
| PGF_04645019 | 17.6 | 310 | 1 | 0 | HPr kinase/phosphorylase |
| PGF_00423732 | 17.59 | 317 | 0.988 | 0.009 | 4-hydroxy-3-methylbut-2-enyl diphosphate reductase (EC 1.17.7.4) |
| PGF_00071514 | 17.53 | 348 | 0.94 | 0.045 | Uncharacterized protein YceB |
| PGF_05165078 | 17.48 | 317 | 0.982 | 0.002 | Methionyl-tRNA formyltransferase (EC 2.1.2.9) |
| PGF_00760084 | 17.45 | 307 | 0.996 | 0 | Cystathionine beta-synthase (EC 4.2.1.22) |
| PGF_04835795 | 17.43 | 304 | 1 | 0 | Site-specific tyrosine recombinase XerC |
| PGF_00012288 | 17.43 | 315 | 0.982 | 0.002 | Homocysteine S-methyltransferase (EC 2.1.1.10) |
| PGF_00017894 | 17.42 | 308 | 0.993 | 0 | LysR-family transcriptional regulator Bsu YtlI |
| PGF_02907590 | 17.4 | 312 | 0.985 | 0.001 | Uncharacterized transporter YcxC, EamA family |

Research supported by the Office of Army Research. The views expressed in this paper are those of the authors and do not reflect the official policy or position of the Department of the Army, Department of Defense, or the U.S. Government.

| | | | | | |
|---|---|---|---|---|---|
| PGF_00064755 | 17.4 | 317 | 0.977 | 0.001 | Uncharacterized iron compound ABC uptake transporter, substrate-binding protein |
| PGF_00063974 | 17.39 | 303 | 0.999 | 0 | UDP-N-acetylenolpyruvoylglucosamine reductase (EC 1.3.1.98) |
| PGF_00058767 | 17.38 | 302 | 1 | 0 | Transcriptional regulator of alpha-acetolactate operon AlsR |
| PGF_00031207 | 17.38 | 305 | 0.995 | 0.001 | Uncharacterized transporter YvbV, EamA family |
| PGF_00064857 | 17.38 | 306 | 0.993 | 0 | Uncharacterized oxidoreductase YqkF |
| PGF_00007027 | 17.34 | 301 | 0.999 | 0 | GTP-binding protein Era |
| PGF_00071153 | 17.34 | 311 | 0.983 | 0 | *Bacillus* subtilis catabolite repression transcription factor CcpB |
| PGF_01496598 | 17.32 | 303 | 0.995 | 0.001 | Chemotaxis protein CheV (EC 2.7.3.-) |
| PGF_02390924 | 17.32 | 318 | 0.971 | 0.022 | 16S rRNA (cytosine(1402)-N(4))-methyltransferase (EC 2.1.1.199) |
| PGF_10569727 | 17.31 | 303 | 0.994 | 0 | LSU rRNA pseudouridine(1911/1915/1917) synthase (EC 5.4.99.23) |
| PGF_02346669 | 17.29 | 335 | 0.945 | 0.047 | Ornithine carbamoyltransferase (EC 2.1.3.3) |
| PGF_03125357 | 17.26 | 304 | 0.99 | 0 | O-Methyltransferase involved in polyketide biosynthesis |
| PGF_00425662 | 17.24 | 299 | 0.997 | 0.001 | FIG00002411: Thioredoxin-fold protein |
| PGF_12669939 | 17.2 | 305 | 0.985 | 0.005 | Ribose ABC transporter, substrate-binding protein RbsB (TC 3.A.1.2.1) |
| PGF_09162729 | 17.2 | 298 | 0.996 | 0 | Flagellar hook-associated protein FlgL |
| PGF_00037828 | 17.17 | 298 | 0.994 | 0 | Protease HtpX |
| PGF_10357700 | 17.16 | 302 | 0.988 | 0 | Similar to ribosomal large subunit pseudouridine synthase D, *Bacillus* subtilis YhcT type |
| PGF_09221959 | 17.16 | 301 | 0.989 | 0 | Methylisocitrate lyase (EC 4.1.3.30) |
| PGF_03889350 | 17.1 | 294 | 0.998 | 0 | Uncharacterized protein YgxA |
| PGF_05797639 | 17.08 | 296 | 0.993 | 0 | Predicted rhamnose oligosaccharide ABC transport system, permease component |
| PGF_00734140 | 17.06 | 297 | 0.99 | 0 | hypothetical protein |
| PGF_03136713 | 17.06 | 292 | 0.998 | 0 | Efflux ABC transporter, ATP-binding protein |
| PGF_00017897 | 17.03 | 293 | 0.995 | 0 | LysR-family transcriptional regulator Bsu YwqM |
| PGF_00549380 | 17 | 290 | 0.999 | 0 | 4-hydroxy-tetrahydrodipicolinate synthase (EC 4.3.3.7) |
| PGF_00017893 | 16.98 | 290 | 0.997 | 0 | LysR-family transcriptional regulator BsdA |
| PGF_00006359 | 16.94 | 299 | 0.98 | 0 | Fructokinase (EC 2.7.1.4) |
| PGF_02920362 | 16.93 | 287 | 0.999 | 0 | Cell wall-binding protein YocH |
| PGF_00057415 | 16.9 | 288 | 0.996 | 0 | Transcription antiterminator GlcT |
| PGF_07528295 | 16.89 | 289 | 0.994 | 0 | hypothetical protein |
| PGF_02904727 | 16.89 | 287 | 0.997 | 0 | hypothetical protein |
| PGF_04047965 | 16.87 | 293 | 0.985 | 0.003 | hypothetical protein |

Research supported by the Office of Army Research. The views expressed in this paper are those of the authors and do not reflect the official policy or position of the Department of the Army, Department of Defense, or the U.S. Government.

| | | | | | |
|---|---|---|---|---|---|
| PGF_00020984 | 16.82 | 283 | 1 | 0 | Methenyltetrahydrofolate cyclohydrolase (EC 3.5.4.9) / Methylenetetrahydrofolate dehydrogenase (NADP+) (EC 1.5.1.5) |
| PGF_00064796 | 16.81 | 294 | 0.981 | 0.013 | Uncharacterized membrane protein Bsu0528 (YdeO) |
| PGF_04872827 | 16.79 | 283 | 0.998 | 0 | Chromosome (plasmid) partitioning protein ParB-2 |
| PGF_05682204 | 16.77 | 282 | 0.999 | 0 | Glycine betaine ABC transport system, permease protein OpuAB |
| PGF_00426701 | 16.74 | 281 | 0.998 | 0 | FIG005590: DegV family protein |
| PGF_02971400 | 16.73 | 280 | 1 | 0 | Uncharacterized oxidoreductase YtbE |
| PGF_00059807 | 16.72 | 283 | 0.994 | 0 | Uncharacterized transcriptional regulator YbbH, RpiR family |
| PGF_03150126 | 16.72 | 285 | 0.99 | 0 | hypothetical protein |
| PGF_00409415 | 16.72 | 282 | 0.996 | 0.003 | UPF0750 membrane protein YvjA |
| PGF_00026864 | 16.72 | 281 | 0.997 | 0 | Octanoyl-[GcvH]:protein N-octanoyltransferase (EC 2.3.1.204) |
| PGF_00426877 | 16.71 | 282 | 0.995 | 0 | LSU ribosomal maturation GTPase RbgA (*B. subtilis* YlqF) |
| PGF_00419444 | 16.68 | 290 | 0.979 | 0 | Conserved protein LiaG in *B. subtilis* in Lia cluster |
| PGF_00047141 | 16.66 | 281 | 0.994 | 0 | Alpha-arabinosides ABC transport system, permease protein AraQ |
| PGF_00049914 | 16.64 | 282 | 0.991 | 0.002 | STAS domain protein |
| PGF_00016393 | 16.64 | 277 | 1 | 0 | LSU ribosomal protein L2p (L8e) |
| PGF_00064875 | 16.64 | 285 | 0.986 | 0.001 | Uncharacterized phosphatase YwpJ |
| PGF_03514578 | 16.63 | 277 | 0.999 | 0 | Beta-glucoside bgl operon antiterminator, BglG family |
| PGF_00019066 | 16.63 | 296 | 0.966 | 0.007 | Maltodextrose utilization protein YvdJ |
| PGF_07735780 | 16.62 | 278 | 0.997 | 0 | hypothetical protein |
| PGF_03466304 | 16.61 | 281 | 0.991 | 0 | Quorum-quenching lactonase YtnP |
| PGF_00026863 | 16.58 | 278 | 0.994 | 0.001 | Octanoate-[acyl-carrier-protein]-protein-N-octanoyltransferase, LipM (EC 2.3.1.181) |
| PGF_03589817 | 16.58 | 282 | 0.987 | 0 | Peptidoglycan N-acetylglucosamine deacetylase (EC 3.5.1.-) |
| PGF_00014397 | 16.53 | 278 | 0.992 | 0 | Inosose isomerase (EC 5.3.99.11) |
| PGF_03152833 | 16.53 | 321 | 0.923 | 0.059 | Ku domain protein |
| PGF_01019139 | 16.52 | 280 | 0.987 | 0.002 | Uncharacterized oxidoreductase YusZ |
| PGF_06874321 | 16.48 | 276 | 0.992 | 0 | Multiple sugar ABC transporter, permease protein MsmG |
| PGF_02475999 | 16.47 | 274 | 0.995 | 0.004 | RsbT co-antagonist protein RsbRA |
| PGF_02868448 | 16.46 | 275 | 0.993 | 0 | Uncharacterized transcriptional regulator YrhO, TrmB family |
| PGF_02781328 | 16.46 | 272 | 0.998 | 0 | NAD synthetase (EC 6.3.1.5) |
| PGF_12908096 | 16.45 | 280 | 0.983 | 0.015 | Spermidine synthase (EC 2.5.1.16) |
| PGF_00026300 | 16.43 | 271 | 0.998 | 0 | Novel pyridoxal kinase, thiD family (EC 2.7.1.35) |
| PGF_12832328 | 16.42 | 278 | 0.985 | 0.002 | BmrR regulator of multidrug efflux transporter Bmr |

Research supported by the Office of Army Research. The views expressed in this paper are those of the authors and do not reflect the official policy or position of the Department of the Army, Department of Defense, or the U.S. Government.

| | | | | | |
|---|---|---|---|---|---|
| PGF_04149001 | 16.4 | 270 | 0.998 | 0 | Hydroxymethylpyrimidine ABC transporter, transmembrane component |
| PGF_00045950 | 16.32 | 272 | 0.99 | 0 | Pyrroline-5-carboxylate reductase (EC 1.5.1.2), ProG-like |
| PGF_03375596 | 16.3 | 270 | 0.992 | 0 | Sugar phosphatase YidA (EC 3.1.3.23) |
| PGF_01176589 | 16.3 | 267 | 0.997 | 0 | 4-hydroxy-tetrahydrodipicolinate reductase (EC 1.17.1.8) |
| PGF_01329082 | 16.3 | 289 | 0.959 | 0.021 | 23S rRNA (guanine(748)-N(1))-methyltransferase (EC 2.1.1.188) |
| PGF_00009882 | 16.29 | 268 | 0.995 | 0 | Uncharacterized glycosyltransferase YwdF |
| PGF_00053997 | 16.29 | 267 | 0.997 | 0.001 | Stage 0 sporulation two-component response regulator (Spo0A) |
| PGF_00026200 | 16.23 | 273 | 0.982 | 0.007 | Non-heme chloroperoxidase (EC 1.11.1.10) |
| PGF_01127158 | 16.23 | 264 | 0.999 | 0 | Zn-dependent hydrolase YycJ/WalJ, required for cell wall metabolism and coordination of cell division with DNA replication |
| PGF_00005450 | 16.22 | 264 | 0.998 | 0 | Flagellar hook protein FlgE |
| PGF_09027360 | 16.19 | 264 | 0.996 | 0 | Regulatory protein RecX |
| PGF_05740384 | 16.16 | 292 | 0.946 | 0.048 | Formamidopyrimidine-DNA glycosylase (EC 3.2.2.23) |
| PGF_03125026 | 16.15 | 271 | 0.981 | 0.005 | Putative uroporphyrinogen-III synthase (EC 4.2.1.75), related to YjjA (in BS) |
| PGF_00083876 | 16.14 | 282 | 0.961 | 0.003 | hypothetical protein |
| PGF_00015827 | 16.09 | 412 | 0.793 | 0.202 | Kef-type K+ transport system, predicted NAD-binding component |
| PGF_00046553 | 16.04 | 260 | 0.995 | 0.002 | RNA polymerase sporulation specific sigma factor SigG |
| PGF_00029831 | 16.03 | 280 | 0.958 | 0.04 | PTS system, fructose-specific IIC component |
| PGF_00024307 | 16.01 | 275 | 0.965 | 0.03 | NAD kinase (EC 2.7.1.23) homolog |
| PGF_05318435 | 16 | 257 | 0.998 | 0 | Electron transfer flavoprotein, beta subunit |
| PGF_00401136 | 16 | 257 | 0.998 | 0 | Uncharacterized membrane protein YceF |
| PGF_00286418 | 15.98 | 258 | 0.995 | 0 | Uncharacterized membrane protein YhfC |
| PGF_00010468 | 15.97 | 257 | 0.996 | 0 | Hydrolase (HAD superfamily) in cluster with DUF1447 |
| PGF_02177520 | 15.97 | 255 | 1 | 0 | RNA polymerase sporulation specific sigma factor SigF |
| PGF_00943546 | 15.96 | 260 | 0.99 | 0 | Methylglutaconyl-CoA hydratase (EC 4.2.1.18) |
| PGF_00876160 | 15.94 | 255 | 0.998 | 0 | Alpha-acetolactate decarboxylase (EC 4.1.1.5) |
| PGF_00419622 | 15.9 | 254 | 0.998 | 0 | Coproheme decarboxylase HemQ (no EC) |
| PGF_00071264 | 15.9 | 253 | 1 | 0 | Bacitracin export ATP-binding protein BceA |
| PGF_08131899 | 15.89 | 256 | 0.993 | 0 | hypothetical protein |
| PGF_00423593 | 15.87 | 259 | 0.986 | 0.007 | Dipeptidyl aminopeptidases/acylaminoacyl-peptidase |
| PGF_03145879 | 15.86 | 279 | 0.95 | 0.037 | L-cystine ABC transporter, substrate-binding protein TcyK |

Research supported by the Office of Army Research. The views expressed in this paper are those of the authors and do not reflect the official policy or position of the Department of the Army, Department of Defense, or the U.S. Government.

| | | | | | |
|---|---|---|---|---|---|
| PGF_07099142 | 15.85 | 256 | 0.991 | 0 | Uncharacterized transporter YwcJ, NirC family |
| PGF_00011536 | 15.84 | 254 | 0.994 | 0.002 | Heptaprenyl diphosphate synthase component I (EC 2.5.1.30) |
| PGF_00046280 | 15.84 | 251 | 1 | 0 | RNA polymerase heat shock sigma factor SigI |
| PGF_00071195 | 15.82 | 255 | 0.991 | 0 | Dihydroanticapsin 7-dehydrogenase (EC 1.1.1.385) => BacC |
| PGF_12786716 | 15.81 | 254 | 0.992 | 0 | Transcriptional regulator GlvR, RpiR family |
| PGF_04710902 | 15.81 | 260 | 0.98 | 0.012 | Lactam utilization protein LamB |
| PGF_00391863 | 15.77 | 253 | 0.992 | 0 | Uncharacterized protein YtbQ |
| PGF_00060533 | 15.75 | 254 | 0.989 | 0 | Transmembrane component YkoC of energizing module of thiamin-regulated ECF transporter for HydroxyMethylPyrimidine |
| PGF_05689837 | 15.73 | 250 | 0.995 | 0 | Probable iron export permease protein FetB |
| PGF_02144826 | 15.73 | 249 | 0.997 | 0 | FMN reductase [NAD(P)H] (EC 1.5.1.39) |
| PGF_03888286 | 15.73 | 249 | 0.997 | 0 | FMN reductase (NADPH) (EC 1.5.1.38) |
| PGF_00051581 | 15.73 | 252 | 0.991 | 0 | Short-chain dehydrodenase (gene dltE) |
| PGF_03255161 | 15.73 | 248 | 0.999 | 0 | Tyrosine-protein kinase transmembrane modulator EpsC |
| PGF_00209115 | 15.71 | 250 | 0.993 | 0 | hypothetical protein |
| PGF_12663506 | 15.7 | 254 | 0.985 | 0 | hypothetical protein |
| PGF_08789310 | 15.67 | 247 | 0.997 | 0 | L-cystine ABC transporter, ATP-binding protein TcyC |
| PGF_03506091 | 15.61 | 245 | 0.997 | 0 | UPF0721 transmembrane protein YdhB |
| PGF_00048586 | 15.59 | 245 | 0.996 | 0 | Ribonuclease PH (EC 2.7.7.56) |
| PGF_02999829 | 15.58 | 253 | 0.979 | 0.012 | Uncharacterized protein YvpB |
| PGF_00413208 | 15.53 | 243 | 0.996 | 0 | tRNA (guanine(37)-N(1))-methyltransferase (EC 2.1.1.228) |
| PGF_00761454 | 15.52 | 243 | 0.996 | 0 | hypothetical protein |
| PGF_00058741 | 15.52 | 241 | 1 | 0 | Transcriptional regulator in cluster with Zn-dependent hydrolase |
| PGF_00062143 | 15.45 | 242 | 0.993 | 0.001 | Twin-arginine translocation protein TatCd |
| PGF_00064474 | 15.44 | 240 | 0.997 | 0 | Arginine transport ATP-binding protein ArtM |
| PGF_00020322 | 15.35 | 261 | 0.95 | 0.039 | Metal-binding protein ZinT |
| PGF_07474537 | 15.3 | 234 | 1 | 0 | L-cystine ABC transporter, permease protein TcyB |
| PGF_00015991 | 15.22 | 235 | 0.993 | 0 | L-cystine ABC transporter, permease protein TcyM |
| PGF_00378256 | 15.22 | 245 | 0.972 | 0.013 | Uncharacterized membrane anchored protein YpbE |
| PGF_00805747 | 15.18 | 231 | 0.998 | 0 | ABC transporter ATP-binding protein YtrE |
| PGF_00060190 | 15.1 | 229 | 0.998 | 0 | ABC transporter-like sensor linked response regulator YxdJ |
| PGF_00066735 | 15.09 | 245 | 0.964 | 0.017 | Vng1746c |
| PGF_10525342 | 15.04 | 247 | 0.957 | 0.03 | Transcriptional regulator, AraC family |
| PGF_03017026 | 15.02 | 227 | 0.997 | 0 | hypothetical protein |

Research supported by the Office of Army Research. The views expressed in this paper are those of the authors and do not reflect the official policy or position of the Department of the Army, Department of Defense, or the U.S. Government.

| | | | | | |
|---|---|---|---|---|---|
| PGF_07723617 | 15 | 225 | 1 | 0 | Putative membrane protease YugP |
| PGF_02992152 | 14.93 | 229 | 0.987 | 0.004 | Uncharacterized protein YcbO |
| PGF_03044001 | 14.92 | 226 | 0.992 | 0 | Tricarboxylate transport transcriptional regulator TctD => Mg-citrate response regulator CitT |
| PGF_03066301 | 14.91 | 240 | 0.963 | 0.029 | Uncharacterized protein YobT |
| PGF_00051120 | 14.91 | 229 | 0.985 | 0.013 | Septum site-determining protein MinC |
| PGF_00065865 | 14.91 | 230 | 0.983 | 0.003 | Uncharacterized transcriptional response regulator YcbL |
| PGF_00317190 | 14.88 | 226 | 0.99 | 0.001 | Teichuronic acid biosynthesis protein TuaF |
| PGF_00047734 | 14.85 | 223 | 0.994 | 0 | Respiratory nitrate reductase gamma chain (EC 1.7.99.4) |
| PGF_00062557 | 14.79 | 235 | 0.965 | 0.022 | Two-component response regulator, malate (EC 2.7.3.-) |
| PGF_00063451 | 14.78 | 221 | 0.994 | 0 | Type III effector HrpW, hairpin with pectate lyase domain |
| PGF_00424244 | 14.78 | 265 | 0.908 | 0.086 | Endo-beta-1,3-1,4 glucanase (licheninase) (EC 3.2.1.73) |
| PGF_06326847 | 14.76 | 221 | 0.993 | 0.004 | Flagellar biosynthesis protein FliP |
| PGF_00052541 | 14.75 | 227 | 0.979 | 0.003 | Amino acid racemase RacX |
| PGF_00425679 | 14.73 | 217 | 1 | 0 | FIG000605: protein co-occurring with transport systems (COG1739) |
| PGF_00038920 | 14.73 | 220 | 0.993 | 0 | Pseudouridine-5' phosphatase (EC 3.1.3.-) |
| PGF_00046555 | 14.71 | 218 | 0.996 | 0.002 | RNA polymerase sporulation specific sigma factor SigH |
| PGF_00067178 | 14.7 | 222 | 0.986 | 0 | Uncharacterized membrane protein YteU |
| PGF_00054072 | 14.69 | 218 | 0.995 | 0.001 | Stage III sporulation protein AH |
| PGF_07320841 | 14.69 | 218 | 0.995 | 0 | ClpCP protease substrate adapter protein MecA |
| PGF_07459774 | 14.65 | 216 | 0.997 | 0 | Succinyl-CoA:3-ketoacid-coenzyme A transferase subunit B (EC 2.8.3.5) |
| PGF_00118588 | 14.65 | 220 | 0.987 | 0.009 | Protease PrsW |
| PGF_00011470 | 14.59 | 213 | 1 | 0 | Hemolysin III |
| PGF_00056362 | 14.57 | 216 | 0.992 | 0 | TPR repeat-containing protein YrrB |
| PGF_03063210 | 14.54 | 213 | 0.996 | 0 | TVP38/TMEM64 family membrane protein YtxB |
| PGF_01776925 | 14.52 | 211 | 0.999 | 0 | Uridine kinase (EC 2.7.1.48) |
| PGF_08089396 | 14.5 | 218 | 0.982 | 0 | Tricarboxylate transport transcriptional regulator TctD |
| PGF_00277008 | 14.46 | 210 | 0.998 | 0 | Uncharacterized membrane protein YrhP |
| PGF_01921960 | 14.46 | 214 | 0.988 | 0 | Uncharacterized sugar epimerase YhfK |
| PGF_00016431 | 14.46 | 209 | 1 | 0 | LSU ribosomal protein L3p (L3e) |
| PGF_00346814 | 14.44 | 212 | 0.992 | 0 | DUF1054 superfamily protein |
| PGF_04861087 | 14.44 | 249 | 0.915 | 0.071 | Two-component response regulator BceR |
| PGF_04867467 | 14.41 | 212 | 0.989 | 0 | Imidazole glycerol phosphate synthase amidotransferase subunit HisH |
| PGF_02445053 | 14.38 | 208 | 0.997 | 0 | Uncharacterized protein YkvT, NOT involved in spore germination |

Research supported by the Office of Army Research. The views expressed in this paper are those of the authors and do not reflect the official policy or position of the Department of the Army, Department of Defense, or the U.S. Government.

| | | | | | |
|---|---|---|---|---|---|
| PGF_03810679 | 14.32 | 205 | 1 | 0 | SOS-response repressor and protease LexA (EC 3.4.21.88) |
| PGF_01756668 | 14.32 | 205 | 1 | 0 | UPF0111 protein YkaA, likely to be a phosphate transport regulator |
| PGF_00027684 | 14.32 | 211 | 0.986 | 0.006 | Acetyltransferase AcuA, acetyl-CoA synthetase inhibitor |
| PGF_00819204 | 14.3 | 221 | 0.962 | 0.021 | Uncharacterized DedA family protein YkoX |
| PGF_00015822 | 14.3 | 205 | 0.999 | 0 | KapD, inhibitor of KinA pathway to sporulation |
| PGF_10458966 | 14.23 | 209 | 0.984 | 0.006 | hypothetical protein |
| PGF_00057526 | 14.19 | 210 | 0.979 | 0.011 | Anti-sigma-W factor RsiW |
| PGF_04192284 | 14.19 | 204 | 0.993 | 0 | Conserved membrane protein in copper uptake, YcnI |
| PGF_03299849 | 14.19 | 204 | 0.993 | 0.001 | Hydroxyaromatic non-oxidative decarboxylase protein B (EC 4.1.1.-) |
| PGF_00049893 | 14.14 | 200 | 1 | 0 | SSU ribosomal protein S4p (S9e) @ SSU ribosomal protein S4p (S9e), zinc-independent |
| PGF_10537005 | 14.12 | 218 | 0.957 | 0.032 | Probable metallo-hydrolase YqgX |
| PGF_00730659 | 14.08 | 200 | 0.995 | 0 | Uncharacterized transcriptional response regulator YvfU |
| PGF_02944297 | 14.06 | 198 | 0.999 | 0 | hypothetical protein |
| PGF_00060189 | 14.05 | 288 | 0.828 | 0.142 | Transcriptional regulatory protein YvrH |
| PGF_00924363 | 14.04 | 199 | 0.995 | 0 | Substrate-specific component YkoE of thiamin-regulated ECF transporter for HydroxyMethylPyrimidine |
| PGF_00285620 | 14.03 | 197 | 1 | 0 | ATP-dependent Clp protease proteolytic subunit ClpP (EC 3.4.21.92) |
| PGF_06649360 | 14.02 | 197 | 0.999 | 0 | Segregation and condensation protein B |
| PGF_06671388 | 14.01 | 200 | 0.991 | 0 | Carboxylesterase (EC 3.1.1.1) |
| PGF_00053894 | 14 | 196 | 1 | 0 | Spore maturation protein A |
| PGF_00039048 | 13.98 | 196 | 0.999 | 0 | Putative 3-methyladenine DNA glycosylase |
| PGF_02230794 | 13.98 | 199 | 0.991 | 0.001 | Molybdenum cofactor guanylyltransferase (EC 2.7.7.77) |
| PGF_00045787 | 13.95 | 196 | 0.996 | 0 | Pyridoxal 5'-phosphate synthase (glutamine hydrolyzing), glutaminase subunit (EC 4.3.3.6) |
| PGF_10556767 | 13.95 | 205 | 0.974 | 0 | HD domain protein |
| PGF_00004058 | 13.9 | 194 | 0.998 | 0 | Fatty acid metabolism regulator protein FadR, TetR family |
| PGF_03084240 | 13.9 | 194 | 0.998 | 0 | Uncharacterized transcriptional regulator YvdT, TetR family |
| PGF_01323228 | 13.89 | 193 | 1 | 0 | Acyl-phosphate:glycerol-3-phosphate O-acyltransferase PlsY (EC 2.3.1.n3) |
| PGF_06195014 | 13.88 | 194 | 0.997 | 0 | Putative nitroreductase family protein SACOL0874 |
| PGF_00046367 | 13.87 | 194 | 0.996 | 0.001 | RNA polymerase sigma factor SigX |
| PGF_00420679 | 13.86 | 200 | 0.98 | 0 | Cytoplasmic thiamin-binding component of thiamin ABC transporter, COG0011 family |

Research supported by the Office of Army Research. The views expressed in this paper are those of the authors and do not reflect the official policy or position of the Department of the Army, Department of Defense, or the U.S. Government.

| | | | | | |
|---|---|---|---|---|---|
| PGF_00026362 | 13.85 | 198 | 0.984 | 0.008 | Nucleoside 5-triphosphatase RdgB (dHAPTP, dITP, XTP-specific) (EC 3.6.1.66) |
| PGF_03887528 | 13.82 | 257 | 0.862 | 0.13 | UPF0702 transmembrane protein YdfR |
| PGF_00012577 | 13.8 | 280 | 0.825 | 0.124 | Uncharacterized protein YmaE |
| PGF_02820109 | 13.77 | 190 | 0.999 | 0 | GTP cyclohydrolase I (EC 3.5.4.16) type 1 |
| PGF_03147263 | 13.72 | 192 | 0.991 | 0 | Metal-sulfur cluster biosynthesis proteins YuaD |
| PGF_08843714 | 13.72 | 189 | 0.998 | 0 | Septum formation protein Maf |
| PGF_00222761 | 13.72 | 238 | 0.889 | 0.081 | Uncharacterized protein YhbD |
| PGF_01677463 | 13.63 | 187 | 0.997 | 0 | RNA polymerase sigma factor SigW |
| PGF_03799365 | 13.6 | 185 | 1 | 0 | Translation elongation factor P |
| PGF_07808527 | 13.56 | 185 | 0.997 | 0 | ADP-ribose pyrophosphatase (EC 3.6.1.13) |
| PGF_00013296 | 13.54 | 186 | 0.993 | 0 | Uncharacterized protein YjgD |
| PGF_00232324 | 13.48 | 184 | 0.993 | 0 | Chromosome-anchoring protein RacA |
| PGF_02897110 | 13.45 | 185 | 0.989 | 0 | 6-phospho-3-hexuloisomerase (EC 5.3.1.27) |
| PGF_00060261 | 13.38 | 180 | 0.998 | 0 | Transcriptional repressor for NAD biosynthesis in gram-positives |
| PGF_00016443 | 13.38 | 179 | 1 | 0 | LSU ribosomal protein L5p (L11e) |
| PGF_00016444 | 13.38 | 179 | 1 | 0 | LSU ribosomal protein L6p (L9e) |
| PGF_00047732 | 13.37 | 184 | 0.986 | 0 | Respiratory nitrate reductase delta chain (EC 1.7.99.4) |
| PGF_00777822 | 13.37 | 185 | 0.983 | 0.003 | hypothetical protein |
| PGF_06274006 | 13.34 | 181 | 0.992 | 0 | ATP synthase delta chain (EC 3.6.3.14) |
| PGF_04457297 | 13.33 | 187 | 0.975 | 0 | 5-formyltetrahydrofolate cyclo-ligase (EC 6.3.3.2) |
| PGF_07133621 | 13.33 | 184 | 0.983 | 0.004 | 16S rRNA (guanine(966)-N(2))-methyltransferase (EC 2.1.1.171) |
| PGF_00053897 | 13.31 | 178 | 0.998 | 0 | Spore maturation protein B |
| PGF_04666784 | 13.31 | 183 | 0.984 | 0 | hypothetical protein |
| PGF_02031958 | 13.29 | 177 | 0.999 | 0 | Possible colicin V production protein |
| PGF_00930759 | 13.19 | 176 | 0.994 | 0 | L,D-transpeptidase |
| PGF_00333474 | 13.14 | 181 | 0.976 | 0 | Protein csk22 |
| PGF_06701488 | 13.14 | 249 | 0.832 | 0.163 | RNA-binding protein Jag |
| PGF_00054042 | 13.03 | 171 | 0.997 | 0 | Stage III sporulation protein AB |
| PGF_10499347 | 13.03 | 173 | 0.991 | 0 | Molybdopterin-guanine dinucleotide biosynthesis protein MobB |
| PGF_00204364 | 13.03 | 175 | 0.985 | 0 | Uncharacterized protein YkkA |
| PGF_00025215 | 13.01 | 174 | 0.986 | 0.011 | Acetolactate synthase small subunit (EC 2.2.1.6) |
| PGF_00007481 | 13 | 172 | 0.991 | 0 | ThiJ/PfpI family protein YhbO |
| PGF_00022467 | 12.97 | 170 | 0.994 | 0 | Molybdenum cofactor biosynthesis protein MoaB |
| PGF_10546429 | 12.96 | 170 | 0.994 | 0.001 | ATP synthase F0 sector subunit b (EC 3.6.3.14) |
| PGF_00413533 | 12.89 | 167 | 0.997 | 0 | Thiol peroxidase, Tpx-type (EC 1.11.1.15) |

Research supported by the Office of Army Research. The views expressed in this paper are those of the authors and do not reflect the official policy or position of the Department of the Army, Department of Defense, or the U.S. Government.

| | | | | | |
|---|---|---|---|---|---|
| PGF_03174068 | 12.88 | 189 | 0.937 | 0.062 | Transcription antitermination protein NusG |
| PGF_01213071 | 12.87 | 166 | 0.999 | 0 | LSU ribosomal protein L10p (P0) |
| PGF_04653134 | 12.87 | 748 | 0.47 | 0.271 | CDP-glycerol:poly(glycerophosphate) glycerophosphotransferase (EC 2.7.8.12) |
| PGF_00024855 | 12.84 | 165 | 0.999 | 0 | NADPH-dependent 7-cyano-7-deazaguanine reductase (EC 1.7.1.13) |
| PGF_00002432 | 12.81 | 167 | 0.991 | 0.002 | Sporulation membrane protein YtrI |
| PGF_00185080 | 12.78 | 185 | 0.94 | 0.011 | hypothetical protein |
| PGF_00065803 | 12.78 | 166 | 0.992 | 0 | Uncharacterized protein, homolog of B.subtilis yhgC |
| PGF_00804722 | 12.67 | 165 | 0.987 | 0.002 | Uncharacterized protein YocC |
| PGF_10386969 | 12.65 | 186 | 0.928 | 0.054 | Acyl-CoA hydrolase (EC 3.1.2.20) |
| PGF_01949670 | 12.64 | 163 | 0.99 | 0.006 | PTS system, fructose-specific IIB component (EC 2.7.1.202) |
| PGF_00033019 | 12.63 | 161 | 0.995 | 0 | Phenolic acid decarboxylase (EC 4.1.1.-) |
| PGF_02927704 | 12.61 | 168 | 0.973 | 0 | hypothetical protein |
| PGF_00412811 | 12.59 | 160 | 0.995 | 0 | Spore coat protein CotF |
| PGF_01446424 | 12.53 | 158 | 0.997 | 0 | Na(+) H(+) antiporter subunit E |
| PGF_00035451 | 12.51 | 162 | 0.983 | 0.004 | Precorrin-2 oxidase (EC 1.3.1.76) |
| PGF_03119533 | 12.51 | 158 | 0.995 | 0 | hypothetical protein |
| PGF_00049433 | 12.5 | 157 | 0.998 | 0 | S-ribosylhomocysteine lyase (EC 4.4.1.21) @ Autoinducer-2 production protein LuxS |
| PGF_07311230 | 12.49 | 156 | 1 | 0 | Positive regulator of CheA protein activity (CheW) |
| PGF_01682834 | 12.48 | 156 | 0.999 | 0 | Bacterial ribosome SSU maturation protein RimP |
| PGF_02715281 | 12.46 | 160 | 0.985 | 0.001 | Uncharacterized membrane protein YoaS |
| PGF_00401663 | 12.46 | 157 | 0.994 | 0 | Transcriptional regulator of the azlBCD operon, AsnC family |
| PGF_00169629 | 12.44 | 191 | 0.9 | 0.082 | Uncharacterized membrane protein YuaF |
| PGF_00027979 | 12.42 | 156 | 0.995 | 0 | Uncharacterized N-acetyltransferase YqjY |
| PGF_03889881 | 12.41 | 154 | 1 | 0 | Transcriptional regulator CtsR |
| PGF_00707140 | 12.36 | 159 | 0.98 | 0.004 | Cys-tRNA(Pro) deacylase YbaK |
| PGF_00022550 | 12.35 | 157 | 0.986 | 0.004 | Molybdopterin synthase catalytic subunit MoaE (EC 2.8.1.12) |
| PGF_00151857 | 12.33 | 178 | 0.924 | 0.04 | hypothetical protein |
| PGF_03090398 | 12.31 | 152 | 0.999 | 0 | Uncharacterized protein YtxH |
| PGF_03501056 | 12.26 | 151 | 0.997 | 0 | hypothetical protein |
| PGF_03021263 | 12.22 | 151 | 0.994 | 0 | SepF, FtsZ-interacting protein related to cell division |
| PGF_02156631 | 12.18 | 149 | 0.998 | 0 | Arginine pathway regulatory protein ArgR, repressor of arg regulon |
| PGF_10470343 | 12.18 | 149 | 0.998 | 0 | Ferric uptake regulation protein FUR |
| PGF_01724713 | 12.17 | 149 | 0.997 | 0 | Ribose-5-phosphate isomerase B (EC 5.3.1.6) |
| PGF_10054809 | 12.16 | 148 | 0.999 | 0 | Transamidase GatB domain protein |
| PGF_00797662 | 12.12 | 147 | 1 | 0 | CBS domain-containing protein YkuL |

Research supported by the Office of Army Research. The views expressed in this paper are those of the authors and do not reflect the official policy or position of the Department of the Army, Department of Defense, or the U.S. Government.

| | | | | | |
|---|---|---|---|---|---|
| PGF_00012657 | 12.12 | 147 | 1 | 0 | ACT domain-containing protein |
| PGF_03802161 | 12.1 | 199 | 0.857 | 0.13 | DNA-directed RNA polymerase delta subunit (EC 2.7.7.6) |
| PGF_03147273 | 12.07 | 146 | 0.999 | 0 | Nitrite-sensitive transcriptional repressor NsrR |
| PGF_00362441 | 12.04 | 164 | 0.94 | 0.042 | hypothetical protein |
| PGF_01222424 | 12.04 | 194 | 0.864 | 0.124 | DinB protein |
| PGF_00002428 | 12.01 | 148 | 0.987 | 0 | Uncharacterized protein YpiF |
| PGF_06594013 | 11.96 | 144 | 0.997 | 0.001 | Peptide-methionine (R)-S-oxide reductase MsrB (EC 1.8.4.12) |
| PGF_00183711 | 11.96 | 145 | 0.993 | 0 | hypothetical protein |
| PGF_10545696 | 11.94 | 148 | 0.982 | 0 | hypothetical protein |
| PGF_09755850 | 11.93 | 143 | 0.997 | 0 | Na(+) H(+) antiporter subunit B |
| PGF_03518570 | 11.92 | 148 | 0.98 | 0.01 | Nucleoside diphosphate kinase (EC 2.7.4.6) |
| PGF_01465362 | 11.92 | 142 | 1 | 0 | Mn-dependent transcriptional regulator MntR |
| PGF_07695531 | 11.87 | 141 | 1 | 0 | LSU ribosomal protein L11p (L12e) |
| PGF_00799099 | 11.87 | 141 | 1 | 0 | Uncharacterized transcriptional regulator YpoP, MarR family |
| PGF_01602421 | 11.8 | 140 | 0.997 | 0 | Flagellar basal-body rod modification protein FlgD |
| PGF_03994339 | 11.75 | 161 | 0.926 | 0.053 | hypothetical protein |
| PGF_10488938 | 11.75 | 138 | 1 | 0 | Putative pre-16S rRNA nuclease YqgF |
| PGF_00053855 | 11.67 | 172 | 0.89 | 0.013 | Spore coat protein CotX |
| PGF_00070331 | 11.63 | 174 | 0.882 | 0.067 | Uncharacterized protein YrrD |
| PGF_00020996 | 11.63 | 137 | 0.994 | 0 | Methylglyoxal synthase (EC 4.2.3.3) |
| PGF_00404745 | 11.63 | 139 | 0.986 | 0 | prolyl endopeptidase |
| PGF_09345013 | 11.62 | 139 | 0.986 | 0.004 | Iron-sulfur cluster regulator IscR |
| PGF_00026337 | 11.59 | 136 | 0.994 | 0.003 | Sporulation-specific extracellular nuclease NucB |
| PGF_03074981 | 11.59 | 136 | 0.994 | 0 | hypothetical protein |
| PGF_00007489 | 11.57 | 146 | 0.958 | 0.041 | UPF0478 protein YtxG |
| PGF_01447607 | 11.55 | 135 | 0.994 | 0 | Uncharacterized CoA-binding protein YneT |
| PGF_03147810 | 11.54 | 134 | 0.997 | 0 | Putative oxidoreductase CatD |
| PGF_00175215 | 11.52 | 135 | 0.991 | 0 | hypothetical protein |
| PGF_00195004 | 11.43 | 143 | 0.956 | 0.017 | hypothetical protein |
| PGF_02019584 | 11.42 | 131 | 0.998 | 0 | Regulatory protein Spx |
| PGF_06348445 | 11.42 | 135 | 0.983 | 0.015 | Anti-sigma B factor RsbT |
| PGF_00373500 | 11.4 | 131 | 0.996 | 0 | hypothetical protein |
| PGF_12835789 | 11.31 | 131 | 0.989 | 0.008 | UPF0297 protein YrzL |
| PGF_00012257 | 11.31 | 128 | 1 | 0 | Holin-like protein CidA |
| PGF_09008105 | 11.25 | 127 | 0.998 | 0 | Chorismate mutase II (EC 5.4.99.5) |
| PGF_00013273 | 11.24 | 128 | 0.994 | 0 | Uncharacterized protein YaeR with similarity to glyoxylase family |
| PGF_07192369 | 11.21 | 128 | 0.991 | 0 | Uncharacterized protein YlqD |

Research supported by the Office of Army Research. The views expressed in this paper are those of the authors and do not reflect the official policy or position of the Department of the Army, Department of Defense, or the U.S. Government.

| | | | | | |
|---|---|---|---|---|---|
| PGF_00309250 | 11.15 | 145 | 0.926 | 0.023 | hypothetical protein |
| PGF_00149187 | 11.15 | 137 | 0.953 | 0.036 | hypothetical protein |
| PGF_00424261 | 11.14 | 124 | 1 | 0 | Uncharacterized protein YurQ |
| PGF_00064808 | 11.13 | 215 | 0.759 | 0.188 | Uncharacterized membrane protein YwmF |
| PGF_12850351 | 11.11 | 179 | 0.83 | 0.149 | hypothetical protein |
| PGF_00016445 | 11.09 | 123 | 1 | 0 | LSU ribosomal protein L7p/L12p (P1/P2) |
| PGF_00015820 | 11.01 | 128 | 0.974 | 0 | KapB, lipoprotein required for KinB pathway to sporulation |
| PGF_00016604 | 10.98 | 128 | 0.971 | 0.011 | Lactoylglutathione lyase and related lyases |
| PGF_00035192 | 10.95 | 126 | 0.976 | 0 | Possible glyoxylase family protein (Lactoylglutathione lyase) (EC 4.4.1.5) |

#### Supplementary Figures

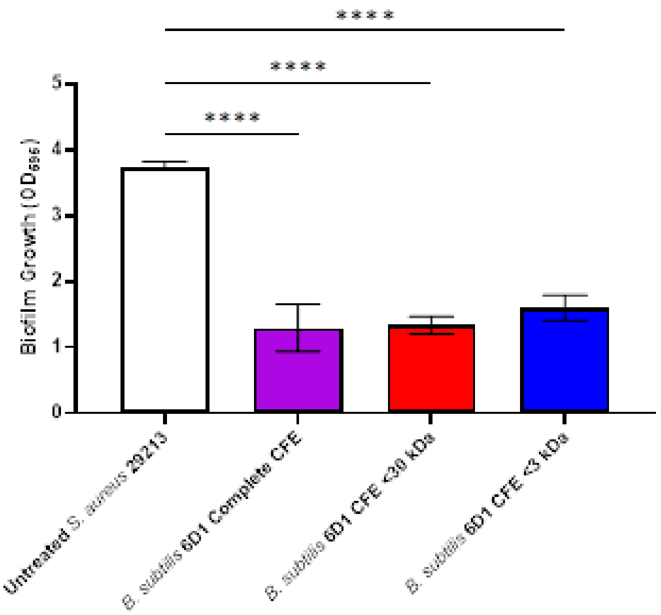


**Figure S1. *B. subtilis* 6D1 CFE harboring low molecular weight metabolites (<3kDa) obtained through centrifugal filtration maintain antibiofilm activity against *S. aureus* ATCC 29213.** *Bacillus* CFE fractions (>30kDa, <30kDa, and <3kDa) obtained via centrifugal filtration were measured for antibiofilm activity to narrow the size range of potentially active compounds. (****) P<0.0001

Research supported by the Office of Army Research. The views expressed in this paper are those of the authors and do not reflect the official policy or position of the Department of the Army, Department of Defense, or the U.S. Government.

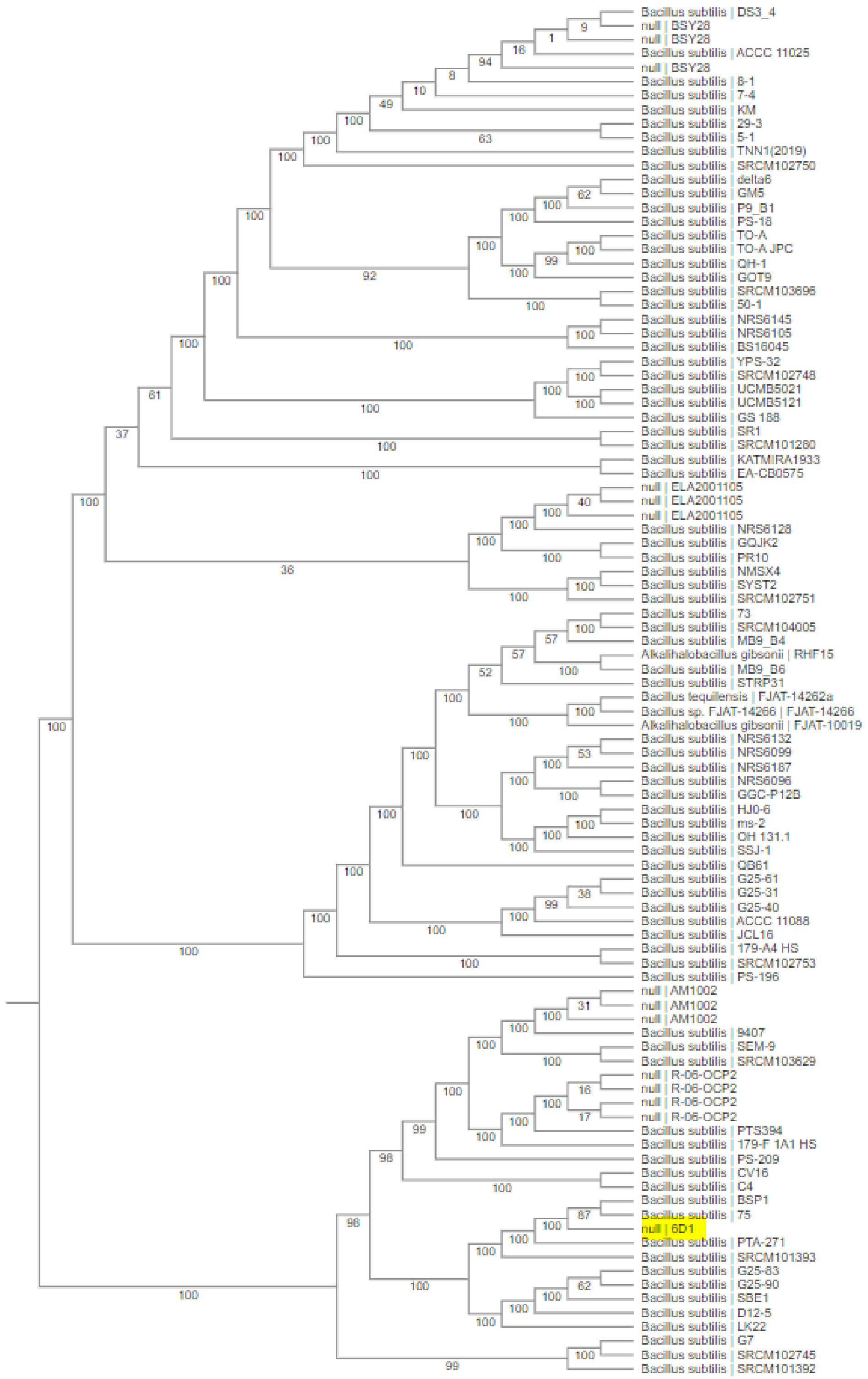


**Figure S2. Phylogenetic analysis comparing *B. subtilis* 6D1 (highlighted) to 98 similarly identified Refseq genomes using 500 random single copy protein coding genes.** *B. subtilis* 75 (Gene Bank Accession # CP045825) and *B. subtilis* PTA-271 (Gene Bank Accession # JACERQ010000010 .1)

Research supported by the Office of Army Research. The views expressed in this paper are those of the authors and do not reflect the official policy or position of the Department of the Army, Department of Defense, or the U.S. Government.

**Figure S3. Biosynthetic gene clusters (BGCs) identified in the *B. subtilis* 6D1 genome.** AntiSmash V6.0 and Clinker were used to visualize and compare reference BGCs (top) to those found in *B. subtilis* 6D1 (bottom).

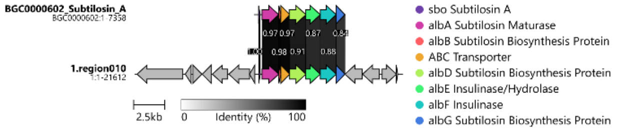


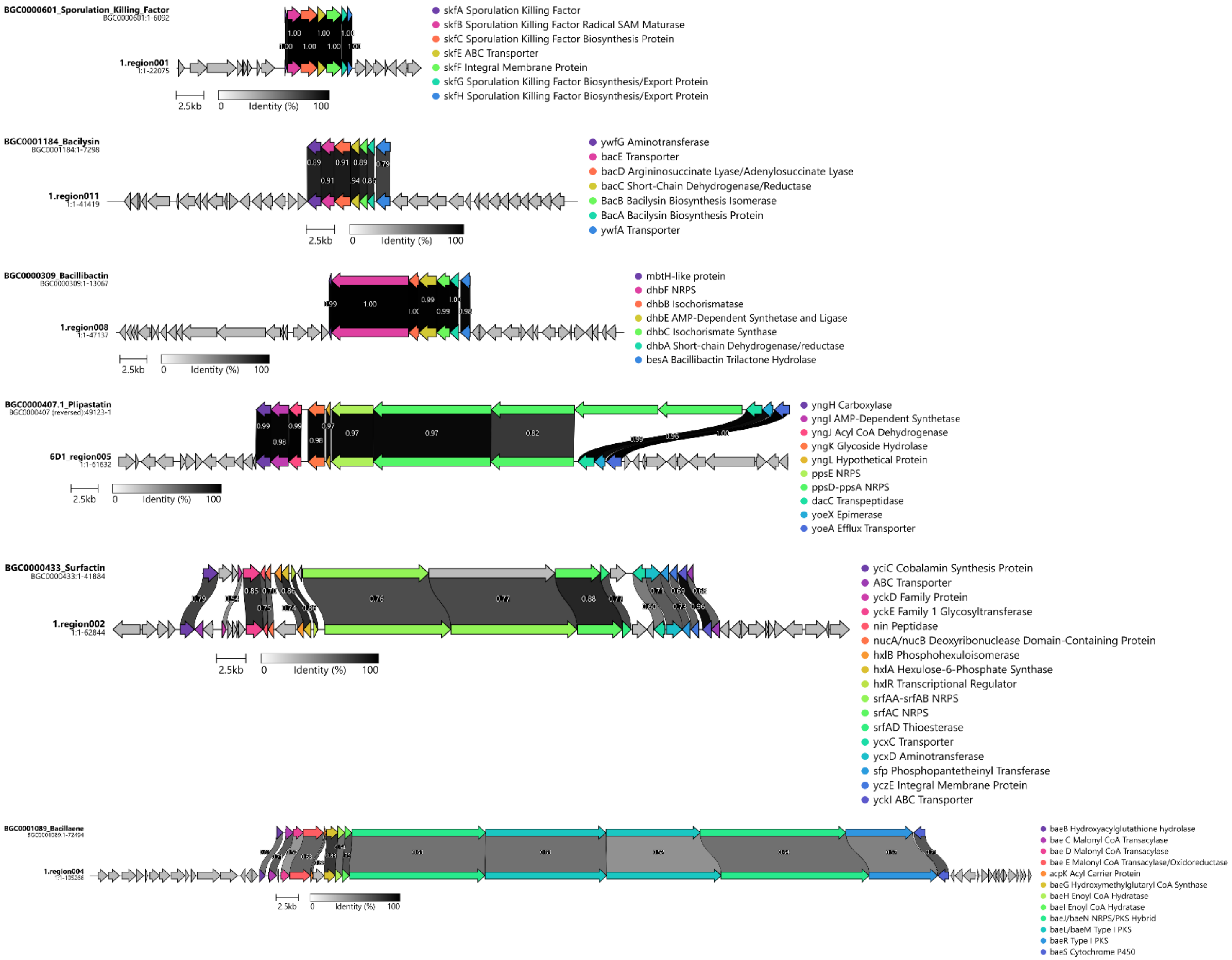


Research supported by the Office of Army Research. The views expressed in this paper are those of the authors and do not reflect the official policy or position of the Department of the Army, Department of Defense, or the U.S. Government.

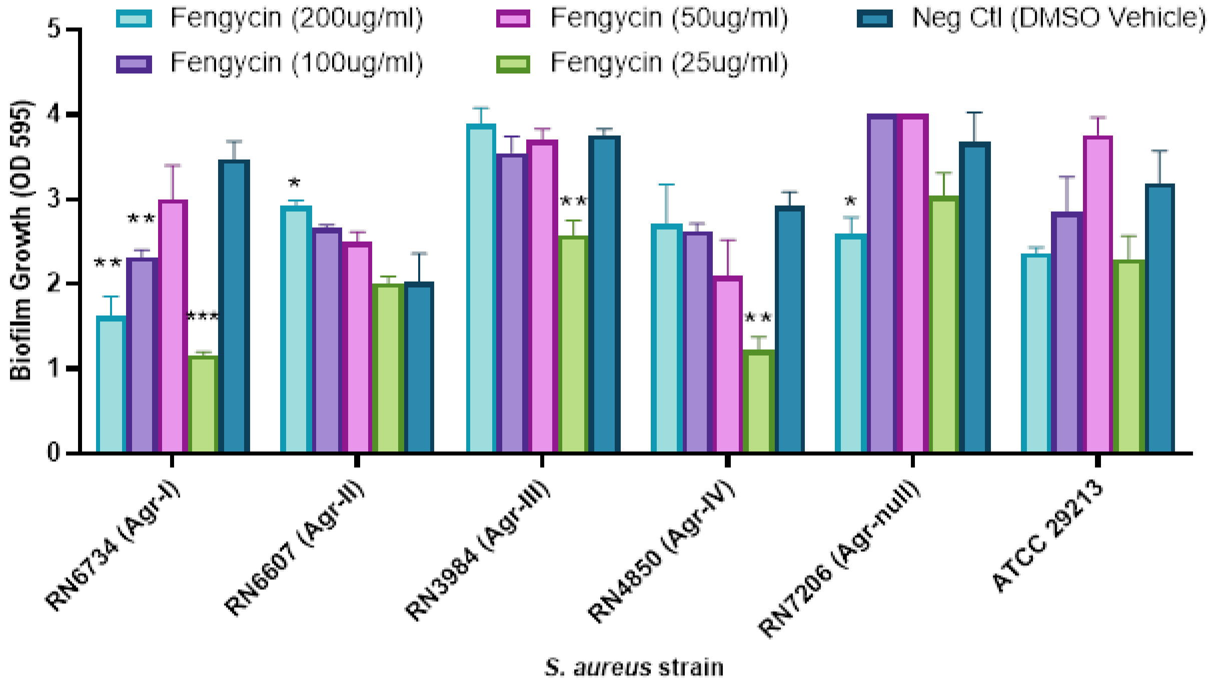


**Figure S4. Commercial HPLC grade fengycin obtained from *B. subtilis* increases *S. aureus* biofilm formation in a concentration and strain specific manner.** (*) P<0.05 (**) P<0.01 (***) P<0.001 compared to the control.

Research supported by the Office of Army Research. The views expressed in this paper are those of the authors and do not reflect the official policy or position of the Department of the Army, Department of Defense, or the U.S. Government.

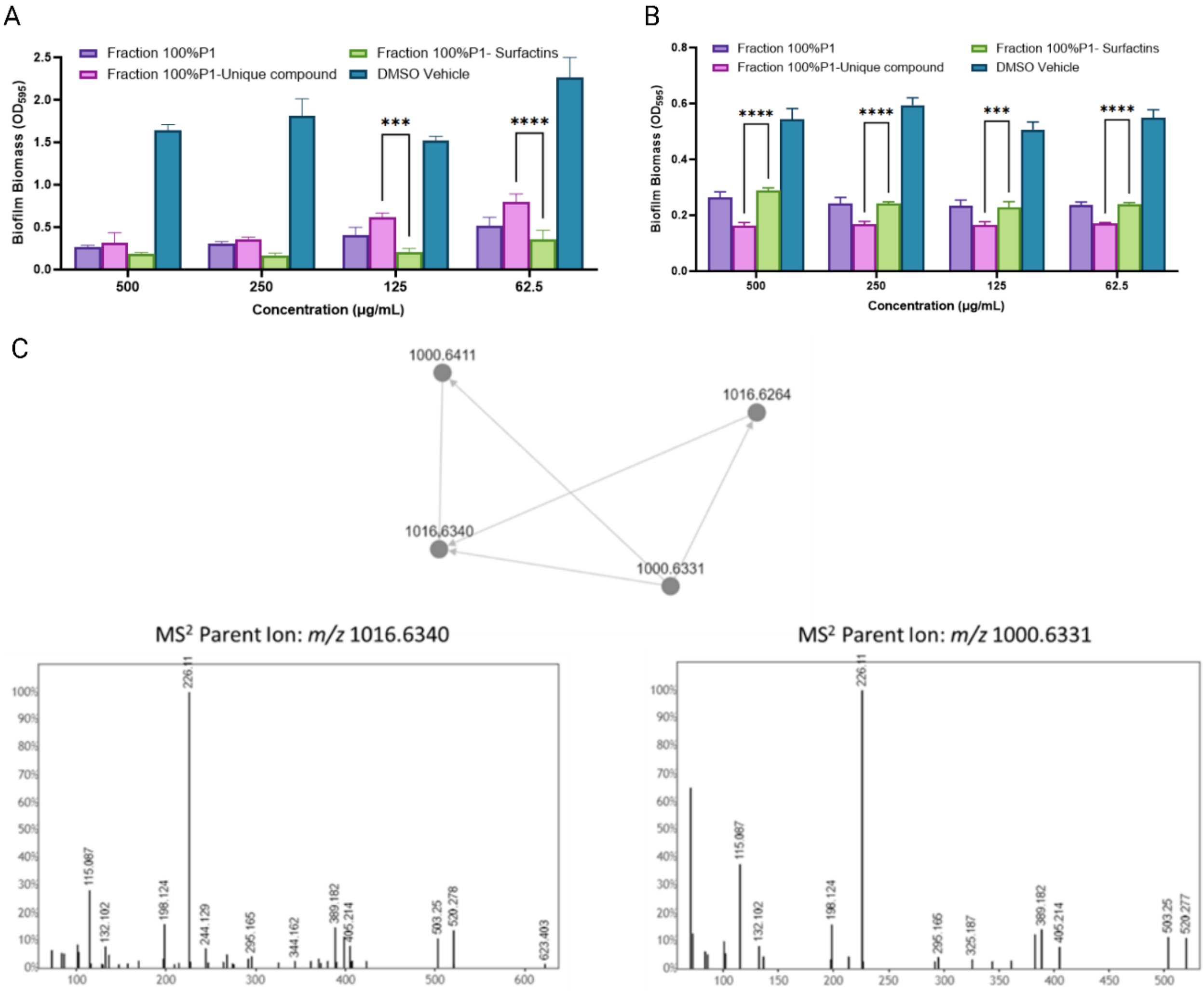


**Figure S5. Unique compound identified in *B. subtilis* 6D1 fraction 100%P1 inhibits *S. aureus* biofilm growth and disrupts mature biofilm.**
**(A)** Biofilm growth inhibition quantified using crystal violet (CV) staining. Either DMSO or *B. subtilis* 6D1 refined fractions were applied to *S. aureus* populations at T0 and incubated statically at 37 ºC for 24 hours. Residual biofilm biomass was quantified by CV staining and averaged across three independent experiments. **(B)** Biofilm disruption quantified using CV straining. Either DMSO or *B. subtilis* 6D1 refined fractions were applied to *S. aureus* 24-hour biofilms and incubated at 37ºC shaking at 100RPM for 2 hours. Residual biofilm biomass was quantified by crystal violet staining and averaged across three independent experiments. Differences were analyzed using a 2-tailed unpaired t-test where (****) $P<0.0001$, (***) $P<0.001$ **(C)** LC-MS/MS spectra and molecular network of *B. subtilis* 6D1 fraction 100%P1 novel compound. Nodes (m/z values) that are connected have a cosine score of at least 0.7 and represent similar molecules. Black spectra represent peaks identified in fraction P1-100% that did not match any known molecules in the GNPS database. The x-axis represents m/z and the y-axis represents the relative abundance of the various ions.

Research supported by the Office of Army Research. The views expressed in this paper are those of the authors and do not reflect the official policy or position of the Department of the Army, Department of Defense, or the U.S. Government.

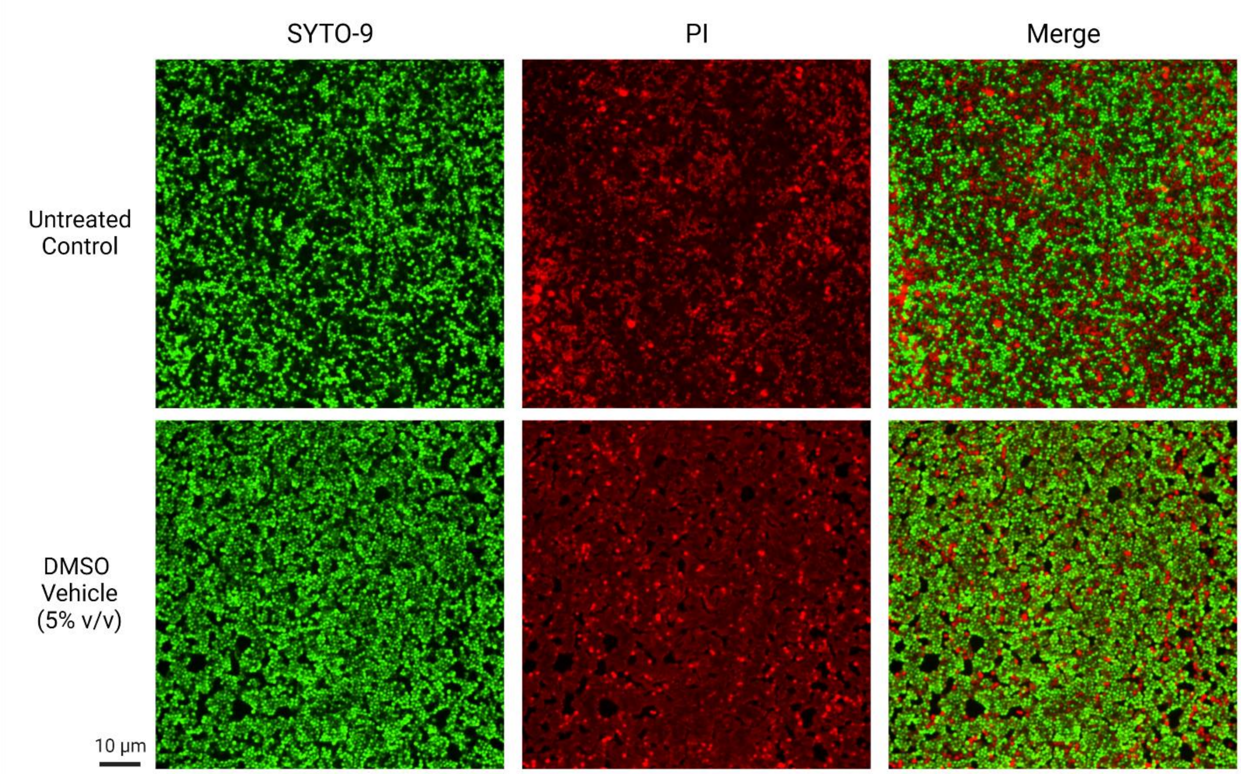


**Figure S6. DMSO applied up to 5% v/v does not alter *S. aureus* ATCC 29213 biofilm growth or increase cell death compared to an untreated control.** Scale bar = 10µm**.**

Research supported by the Office of Army Research. The views expressed in this paper are those of the authors and do not reflect the official policy or position of the Department of the Army, Department of Defense, or the U.S. Government.
